# scTenifoldKnk: an efficient virtual knockout tool for gene function predictions via single-cell gene regulatory network perturbation

**DOI:** 10.1101/2021.03.22.436484

**Authors:** Daniel Osorio, Yan Zhong, Guanxun Li, Qian Xu, Yongjian Yang, Yanan Tian, Robert S. Chapkin, Jianhua Z. Huang, James J. Cai

**Affiliations:** Department of Veterinary Integrative Biosciences, Texas A&M University, College Station, TX 77843, USA; Key Laboratory of Advanced Theory and Application in Statistics and Data Science-MOE, School of Statistics, East China Normal University, Shanghai 200062, China; Department of Statistics, Texas A&M University, College Station, TX 77843, USA; Department of Electrical and Computer Engineering, Texas A&M University, College Station, TX 77843, USA; Department of Veterinary Physiology and Pharmacology, Texas A&M University, College Station, TX 77843, USA; Department of Nutrition, Texas A&M University, College Station, TX 77843, USA; Department of Biochemistry & Biophysics, Texas A&M University, College Station, TX 77843, USA; School of Data Science, The Chinese University of Hong Kong, Shenzhen, Guangdong 518172, China; Interdisciplinary Program of Genetics, Texas A&M University, College Station, TX 77843, USA

**Author notes:** Corresponding authors: J.Z.H. and J.J.C. Lead Contact: J.J.C.

## Abstract

**Bigger Picture:**

- Gene knockout (KO) experiments, using genetically altered animals, are a proven powerful approach to elucidate the role of a gene in a biological process. However, systematic KO experiments targeting many genes are usually prohibitive due to limited experimental and animal resources. Here, we present scTenifoldKnk, an efficient virtual KO tool that allows the systematic deletion of many genes individually. scTenifoldKnk uses single-cell RNA sequencing (scRNAseq) data from wild-type (WT) samples to predict gene function in a cell type-specific manner. We show that predictions made by scTenifoldKnk recapitulate findings from real-animal KO experiments. scTenifoldKnk has proven to be a powerful and effective approach for elucidating gene function, prioritizing KO targets, predicting experimental outcomes before real-animal KO experiments are conducted.

**Highlights:**

- scTenifoldKnk performs virtual KO experiments using scRNAseq data.
- scTenifoldKnk only requires data from WT samples; no data is needed from KO samples.
- Predictions made by scTenifoldKnk recapitulate findings from real-animal KO experiments.

**Data Science Maturity Level:** 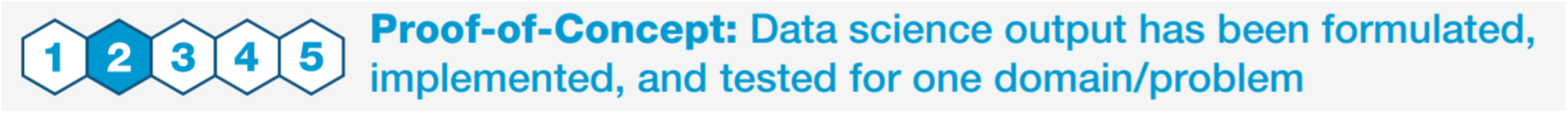

**eTOC blurb:** scTenifoldKnk is a machine learning workflow performing virtual KO experiments to predict gene function. It constructs gene regulatory networks using single-cell RNA sequencing data from wild-type samples and then computationally deletes target genes. Real-data applications demonstrate that scTenifoldKnk recapitulates findings of real-animal KO experiments and accurately predicts gene function in analyzed cells.

**Summary:** Gene knockout (KO) experiments are a proven, powerful approach for studying gene function. However, systematic KO experiments targeting a large number of genes are usually prohibitive due to the limit of experimental and animal resources. Here, we present scTenifoldKnk, an efficient virtual KO tool that enables systematic KO investigation of gene function using data from single-cell RNA sequencing (scRNAseq). In scTenifoldKnk analysis, a gene regulatory network (GRN) is first constructed from scRNAseq data of wild-type samples, and a target gene is then virtually deleted from the constructed GRN. Manifold alignment is used to align the resulting reduced GRN to the original GRN to identify differentially regulated genes, which are used to infer target gene functions in analyzed cells. We demonstrate that the scTenifoldKnk-based virtual KO analysis recapitulates the main findings of real-animal KO experiments and recovers the expected functions of genes in relevant cell types.

## Introduction

Gene knockout (KO) experiments are a proven approach for studying gene function. A typical KO experiment involves the phenotypic characterization of organisms following the deletion of a target gene. For example, in KO mice, a gene is knocked out, i.e., made inoperative by deleting one or more alleles using genetic techniques. Phenotypic characterization of KO animals provides insight into how the target gene functions within the biological context that KO animals present. Notably, gene functions can be inferred by contrasting phenotypes between KO and wild-type (WT) animals and identifying differences. At the molecular level, gene expression may serve as a quantitative phenotype. The state of gene expression is regulated in a coordinated manner in all living organisms, exhibiting synchronized patterns of transcription that can be depicted using a gene regulatory network (GRN). Co-regulations are often seen among genes associated with the same biological processes and pathways or regulated by the same master transcription factors (TFs).^1^ If one gene is knocked out, functionally related genes can mediate a homeostatic response. Thus, in unraveling regulatory mechanisms and synchronized patterns of cellular transcriptional activities, network analysis of gene expression provides mechanistic insights.

The advent of single-cell technology has greatly improved cellular phenotyping resolution. For example, high-throughput droplet-based single-cell RNA sequencing (scRNAseq) makes it possible to profile transcriptomes of thousands of individual cells in a single experiment. The application of single-cell technology in KO experiments allows investigators to probe gene function in a cell type-specific manner. To fully understand the regulatory mechanisms and gene function, it would be ideal for applying scRNAseq in the context of systematic KO experiments that involve the deletion of many genes individually. This unique coupling of scRNAseq with large-scale gene KO in an organism would allow the function of many genes in various cell types to be studied at a single-cell level of resolution. However, limited resources tend to prohibit this kind of experiment, especially when multiple genes are targeted and multiple tissues and cell types are involved. To this end, Perturb-seq, and other conceptually similar protocols,^2,3^ have been developed to achieve the aforementioned goal. Although these protocols may allow the study of gene functions in many cells in a massively parallel fashion, they require the creation of large-scale CRISPR libraries,^4^ which presents a major technical challenge. For these reasons, computational methods serve as a possible solution for prohibitive, systematic KO experiments.^5–7^ We propose that new computational methods may take advantage of the synchronized expression of genes in given samples to construct GRNs and predict gene functions. The topology of those GRNs is known to serve as a basis to accurately predict perturbations caused by gene KO.^8^

Here we present a machine learning workflow, scTenifoldKnk, that can be used to perform virtual KO experiments to predict gene functions. scTenifoldKnk utilizes expression data from scRNAseq of the WT samples as input and constructs a denoised single-cell GRN (scGRN). The WT scGRN is copied and then converted to a pseudo-KO scGRN by artificially zeroing out the weight of outward edges of the target gene in the adjacency matrix. Next, by comparing the two scGRNs (WT vs. pseudo-KO), scTenifoldKnk reveals changes in transcriptional regulatory programs and assesses the impact of KO on the WT scGRN. This information is then used to elucidate the functions of the KO gene in analyzed cells through enrichment analysis.

ScTenifoldKnk is computationally efficient enough to allow the method to be applied to systematic KO experiments. In such a systematic study, we assume that thousands of genes in analyzed cells will be knocked out one by one. As mentioned, due to the experimental and biological limitations, such systematic KO experiments would be extremely difficult, if not impossible, to conduct in a real-animal experimental setting. The other application features of scTenifoldKnk include: scTenifoldKnk requires no data from KO samples, as it only utilizes scRNAseq data from WT samples, and scTenifoldKnk can perform multi-gene KO analysis, i.e., knocking out more than one gene at a time.

The remainder of this paper is organized as follows. We first present an overview of the workflow of scTenifoldKnk. We then use simulated data to demonstrate its basic functions, followed by using existing data generated from authentic-animal KO experiments to highlight the use of scTenifoldKnk. These existing data sets contain scRNAseq expression matrices from both WT and KO samples. Although KO data sets were available, they were not used by scTenifoldKnk as input. Instead, the KO data sets were specifically used as positive controls to show that scTenifoldKnk can produce expected results. We also show the use of scTenifoldKnk to reveal the functions of genes underlying three different Mendelian diseases. Finally, we show two cases of systematic KO using scTenifoldKnk.

## Results

### Result 1: The scTenifoldKnk workflow

ScTenifoldKnk takes a single gene-by-cell count matrix from the WT sample as input. The workflow constructs a WT scGRN from the input count matrix and then generates a pseudo-KO scGRN by knocking out a gene from the WT scGRN. Eventually, it employs a network comparison method to compare the pseudo-KO and WT scGRNs to identify differentially regulated (DR) genes. These DR genes are *virtual-KO perturbed genes*—the two terms will be used interchangeably throughout this paper. From the enriched function of these virtual-KO perturbed genes, the function of the KO gene (i.e., the gene that is virtually knocked out) can be inferred. scTenifoldKnk is implemented with a modular structure containing three core modules illustrated in **Figure 1**. The three steps are summarized as follows.

**Figure 1.**
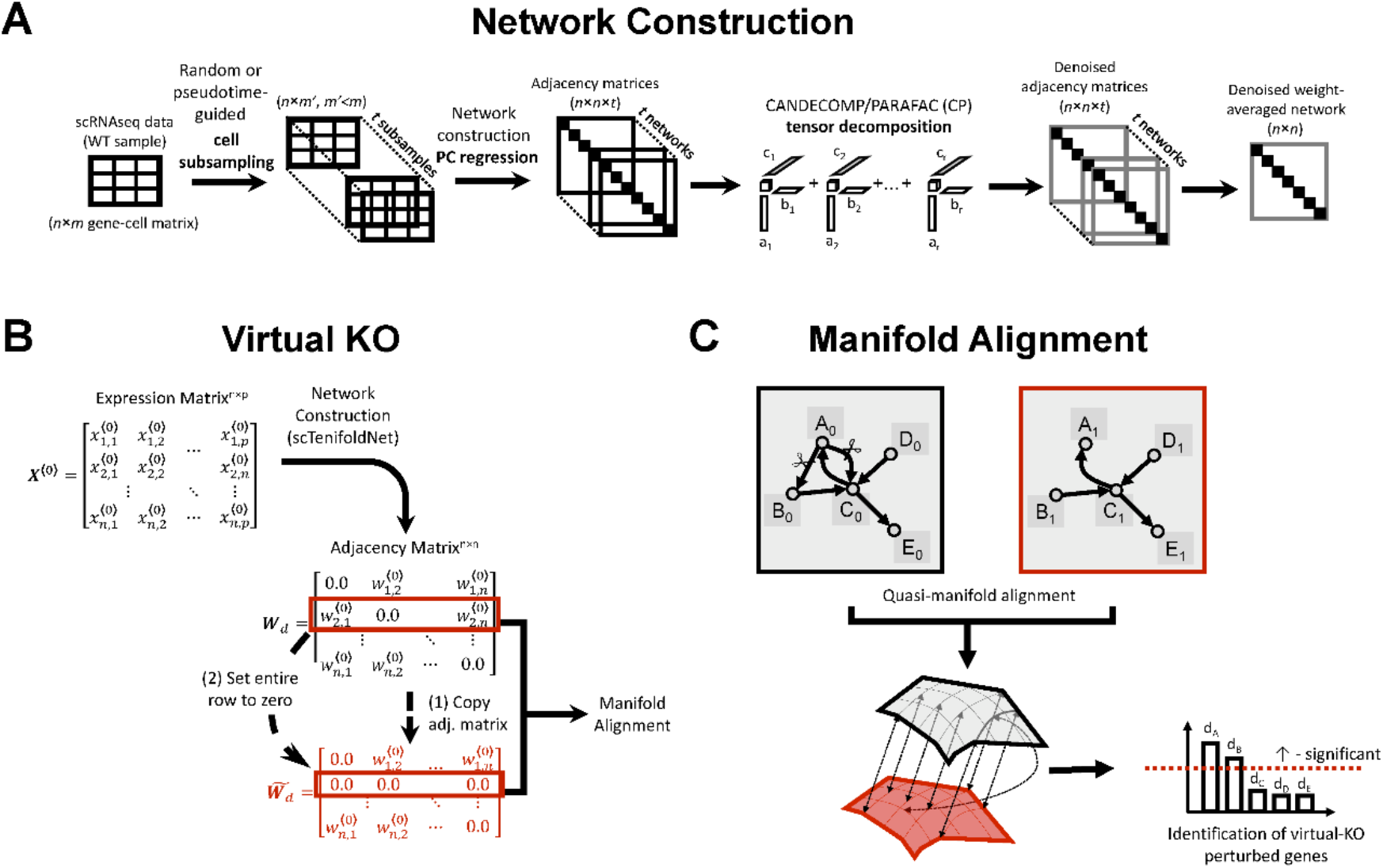
Overview of scTenifoldKnk workflow. ScTenifoldKnk is designed to perform virtual KO experiments with data from scRNAseq. The workflow of scTenifoldKnk consists of three main modules, namely, network construction, virtual KO, and manifold alignment. (**A**) *Network construction*. This module consists of three steps: cell subsampling, principal component regression, and tensor decomposition/denoising. (**B**) *Virtual KO*. This module starts by duplicating the WT adjacency matrix, ***W**_d_*, to make 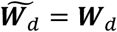. Then, the entire row of 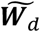 corresponding to the KO gene is set to zero. The modified 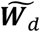 is pseudo-KO scGRN. (**C**) *Manifold alignment*. This method is used to learn latent representations of two networks, ***W**_d_* and 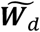, and align them based on their underlying manifold structures. The distance between a gene’s projections with respect to the two scGRNs on the lowdimensional latent representation is used to measure the level of differential regulation of the specific gene. A ranked gene list, in which genes are sorted according to the value of the distance, can be used as input to perform GSEA analysis. The significantly DR genes are identified as *virtual-KO perturbed genes*.

#### Step 1: Constructing scGRN with scRNAseq data from WT samples

With the scRNAseq data from a WT sample, scTenifoldKnk first constructs an scGRN using a pipeline we proposed previously, namely, scTenifoldNet.^9^ This *network construction* step contains three sub-steps (**Figure 1A**):

##### Substep 1.1. Subsampling cells randomly

*Denote **X*** as the scRNAseq expression data matrix, which contains the expression levels for *p* genes and *n* cells. Then, *m* (< n) cells in ***X*** are randomly sampled to form ***X***′ using an *m*-out-of-*n* bootstrap procedure. This subsampling process repeats *t* times to create *t* subsets of cells 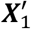,…, 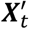.

##### Substep 1.2. Constructing a GRN for each subsampled set of cells

For each 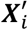, principal component (PC) regression is run *p* times to construct a GRN. Each time the expression level of one gene is used as the response variable and the expression levels for the remaining genes as dependent variables. The constructed GRN from 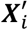 is stored as a signed, weighted and directional graph, represented with a *p* × *p* adjacency matrix ***W**_i_*, each of whose columns stores the regression coefficients for the PC regression of a gene. ***W**_i_* is then normalized via dividing by the maximal absolute value.

##### Substep 1.3. Denoising adjacency matrices to obtain the final GRN

Tensor decomposition is used to denoise the adjacency matrices {***W**_i_*} obtained from the PC regression step. First, the collection of {***W**_i_*} for *t* GRNs is processed as a third-order tensor Ξ, containing *p* × *p* × *t* elements. Next, the CANDECOMP/PARAFAC (CP) decomposition is applied to decompose Ξ into components. Then, Ξ is reconstructed using top *d* components to obtain denoised tensors Ξ_*d*_. Denoised {***W**_i_*} in Ξ_*d*_ are collapsed by taking the average of edge weights to obtain the final averaged matrix, ***W**_i_*.

#### Step 2: Generating pseudo-KO scGRN by virtually knocking out a gene

In the last step, the scRNAseq expression data matrix from the WT sample, ***X***, is first used to construct the WT scGRN. In this step named *virtual KO*, the adjacency matrix of the WT scGRN, ***W**_d_*, is copied, and then the entire row of ***W**_d_* corresponding to the target gene is set to 0 (**Figure 1B**). In this way, the virtual KO operation is performed on ***W**_d_* directly. The modified ***W**_d_* is denoted as 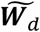, that is, the adjacency matrix of pseudo-KO scGRN.

#### Step 3. Comparing scGRNs to identify virtual-KO perturbed genes

In this step, we assume that WT and pseudo-KO scGRNs, ***W**_d_* and 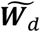, have been obtained. A quasi-manifold alignment method is then used to align ***W**_d_* and 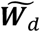 (**Figure 1C**, see **Experimental Procedures** for details). All genes included in the two scGRNs are projected in k-dimensional space, where k << p. After the projection, each gene has two low-dimensional representations: one is in respect to ***W**_d_* and the other 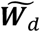. For each gene *j*, *d_j_* is the Euclidean distance between the gene’s two projections. The greater *d_j_*, the more significant the differential regulation. Genes were sorted according to the value of the distance to produce a ranked gene list, which was used as input of the gene set enrichment analysis (GSEA).^10^ Finally, a χ^2^ test is applied to detect significant DR genes, i.e., *virtual-KO perturbed genes*.

A more detailed description of scTenifoldKnk modules is provided in the **Experimental Procedures** section.

### Result 2: Virtual KO analysis using simulated scRNAseq data

We first used the simulated data to validate the relevance of our method. For this purpose, we generated a synthetic scRNAseq data set using the simulator SERGIO—a single-cell expression simulator guided by GRNs.^11^ To simulate the data, we supplied SERGIO with five predefined GRNs of different sizes, containing 5, 10, 25, 40, and 20 genes, respectively. The simulated scRNAseq data set was a sparse matrix (70% zeros) of 3,000 cells and 100 genes. We applied the PC regression method to the simulated data and constructed an scGRN. As expected, in the scGRN, genes were clearly clustered into five distinct modules (see **Figure 2A** for the adjacency matrix), mirroring the predefined modules given by the generative model of SERGIO. Genes in the same module were supposed to be functionally related or under the same regulation. Because SERGIO simulates gene expression in steady-state cells, we only used it to simulate co-expression modules rather than the regulatory processes of some upstream regulators acting on these modules. We regarded the constructed network as the WT scGRN. Next, we set the weight of the 20^th^ gene (gene #20) in the WT scGRN to zero to produce the pseudo-KO scGRN. Gene #20 belongs to the third module, which includes a total of 25 genes. We then used scTenifoldKnk to compare the pseudo-KO scGRN with the WT scGRN to identify genes significantly differentially regulated due to the KO. These genes were predicted likely to be perturbed along with gene #20. Because we knew the KO effect was due to the deletion of gene #20, we expected that the identified genes to be those closely correlated with gene #20. Indeed, as expected, scTenifoldKnk showed all significant genes were from the third module (**Figure 2B & 2C**, top), in which gene #20 is located. We repeated the analysis using genes #50 and #100 as two additional examples. Again, the results were as expected (**Figure 2B & 2C**, middle and bottom). Thus, we concluded that when a member gene is knocked out from a tightly regulated module, other member genes in the same module should be detected by scTenifoldKnk. Algorithmically, genes in the same module with the KO gene were detected because their projected positions in low-dimensional latent representations of WT and pseudo-KO networks changed more than other genes not in the same module (see **Experimental Procedures** for details).

**Figure 2.**
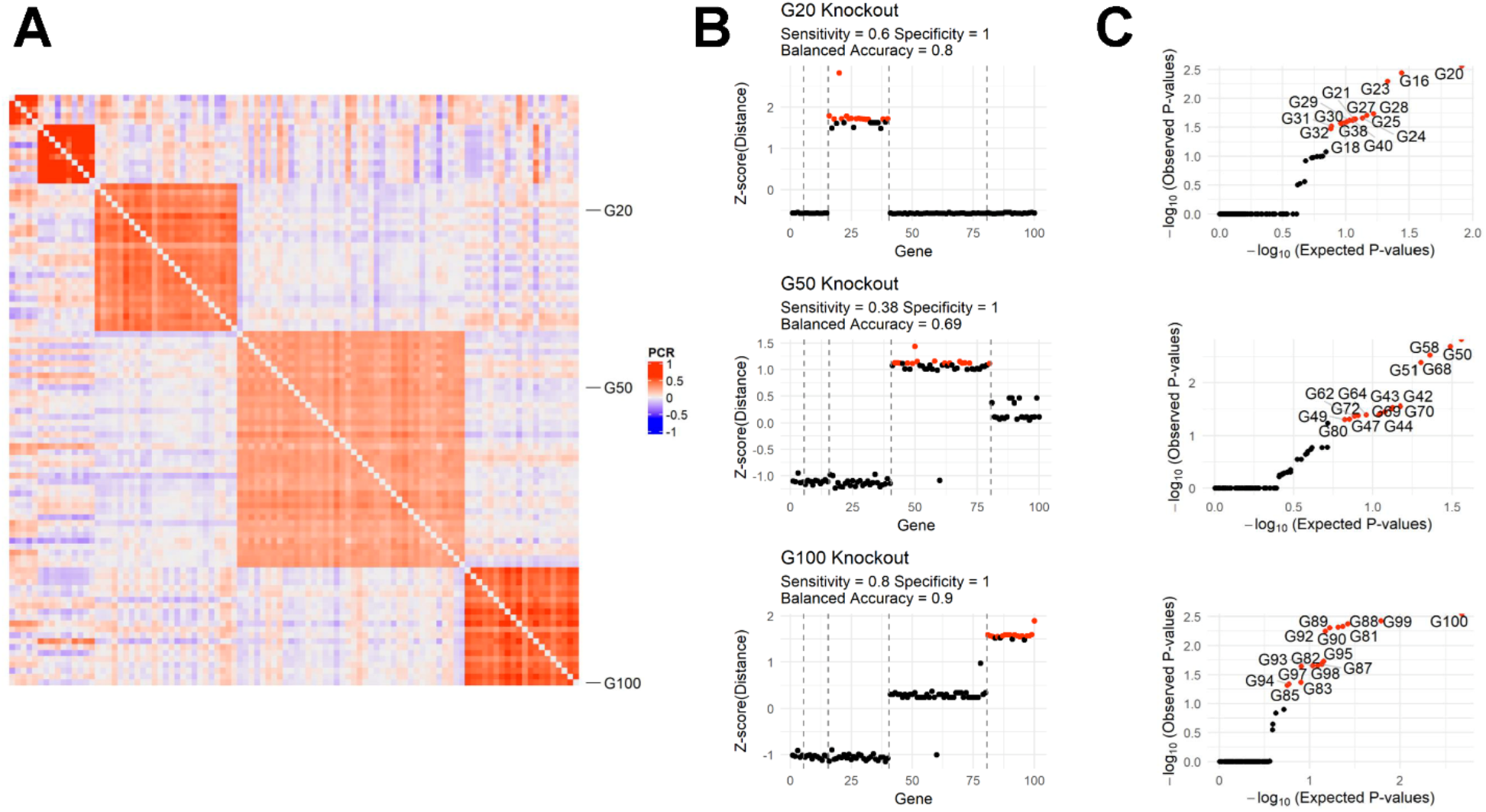
Simulations show that scTenifoldKnk specifically detects regulatory modules that include the KO gene. (**A**) Heatmap of a 100 × 100 adjacency matrix of scGRN constructed from simulated scRNAseq data of 100 genes and 3,000 cells. The color is scaled according to the normalized PC regression coefficient values between gene pairs. The network contains five predefined co-regulated modules of different sizes, indicated by the blocks of gene pairs with a high correlation. The number of genes of each module is 5, 10, 25, 40, and 20, respectively. (**B**) Normalized (Box-Cox) and standardized (Z-score) distance was measured for each gene after manifold alignment and their associated −log10(p-value) after DR testing in the simulated network. Red and black dots indicate whether genes are significant or not after false discovery correction. Assuming genes in the same module should be detected as significant genes, sensitivity is defined as = TP/(TP+FN), and specificity as = (TN)/(TN+FP), where T, P, F, and N stands for true, positive, false, and negative, respectively. Balanced accuracy is defined as the average sensitivity and specificity values. (**C**) QQ-plots of expected (under the uniform distribution) vs. observed p-values of genes given by the DR tests.

### Result 3: scTenifoldKnk virtual KO analysis recapitulates results of real KO experiments

As a virtual KO tool, scTenifoldKnk is expected to recapitulate results obtained from real KO experiments. To prove this, we applied scTenifoldKnk to scRNAseq data from three *in vivo* KO experiments. In all three cases, the scRNAseq data sets of the original studies contained expression matrices from both WT and KO samples.

It is noteworthy that comparisons between predictions made by scTenifoldKnk and the main findings of original papers were by no means “fair” comparisons. This is because scTenifoldKnk was blinded to all information except the scRNAseq data from the WT samples. In contrast, to characterize functions of target genes, original *in vivo* KO studies used scRNAseq data from both WT and KO samples, as well as other empirical data from bulk RNAseq, flow cytometry, and immunostaining assays. Nevertheless, we obtained an overall consistency in the KO gene functions between those predicted using scTenifoldKnk and those reported in the original papers. **Supplementary Table S1** gives a summary of comparisons.

#### Virtual vs. real KO experiment 1: Nkx2-1 is required for the transcriptional control, development, and maintenance of alveolar type-1 and type-2 cells

NK homeobox 2-1 (*Nkx2-1*) is highly expressed in lung epithelial cells of alveolar type I (AT1) and type II (AT2). AT1 cells cover 95% of the gas exchange surface and are 0.1 μm thick to allow passive oxygen diffusion into the bloodstream. *Nkx2-1* is essential at all developmental stages of AT1 cells. Loss of *Nkx2-1* results in the impairment of three main defining features of AT1 cells, molecular markers, expansive morphology, and cellular quiescence.^12^ AT2 cells are cuboidal and secrete surfactants to reduce surface tension. Mutations in *Nkx2-1* interrupt the expression of *Sftpb* and *Sftpc*, two genes related to AT2 cell function and molecular identity.^12,13^

To examine the molecular and cellular changes caused by the *Nkx2-1*^-/-^ KO, Little et al.^12^ generated a comprehensive set of data (GEO: GSM3716703) using the lung samples from WT and *Nkx2-1*^-/-^ KO mice. Using bulk RNAseq and immunostaining assays, they observed that the expression of marker genes of AT1 and AT2 cells was downregulated in the *Nkx2-1* mutant cells. They also found that the expression of marker genes for gastrointestinal cells was upregulated in *Nkx2-1*^-/-^ mutant AT1 cells, which form dense microvilli-like structures apically. Using ChIP-seq, they found that *Nkx2-1* binds to a set of genes implicated in regulating the cytoskeleton, membrane composition, and extracellular matrix. Little et al. also generated scRNAseq data for 2,312 and 2,558 epithelial cells from lung samples of the WT and *Nkx2-1*^-/-^ KO mice, respectively.^12^

We obtained the scRNAseq data, generated by Little et al.,^12^ and used the expression matrix of 8,647 genes × 2,312 cells from WT mice as the input for scTenifoldKnk. We constructed the WT scGRN and then knocked out *Nkx2-1*. The final report of scTenifoldKnk analysis contained 171 significant genes [False Discovery Rate (FDR) < 0.05, **Supplementary Table S2**]. These virtual-KO perturbed genes included 7 out of 32 marker genes of AT1 cells (*Egfl6*, *Ager*, *Cldn18*, *Icam1*, *Crlf1*, *Gprc5a*, and *Aqp5*) and 25 out of 38 marker genes of AT2 cells (highlighted in **Supplementary Table S2**). The functional enrichment test, Enrichr,^14^ indicated that these genes were enriched for functional categories: *epithelial to mesenchymal transition led by the WNT signaling pathway members*, *surfactant homeostasis, lamellar body*, and *cell adhesion molecules*. These enriched functions were consistent with and related to the functions of AT2 cells. Next, we applied the interaction enrichment analysis to the 171 significant genes. The interaction enrichment analysis was provided and based on the STRING protein-protein interaction database.^15^ We found that these genes appear in a fully connected component in the STRING interaction network (p-value < 0.01, STRING interaction enrichment test), indicating a closely related functional relationship between those genes. We subsequently performed GSEA analysis,^10^ to evaluate the extent of perturbation caused by the *Nkx2-1* KO at the transcriptome-wide level. GSEA analysis identified gene sets containing marker genes of AT1 and AT2 cells (FDR < 0.01 in both cases). Specifically, AT1 and AT2 marker genes were among the topmost perturbed genes caused by the deletion of *Nkx2-1*. GSEA analysis also showed that the *Nkx2-1*^-/-^ KO impacted genes with functions related to intestinal microvilli (**Figure 3A**), cell cycle, and the cytoskeleton. These results are consistent with those reported in the original study.^12^

**Figure 3.**
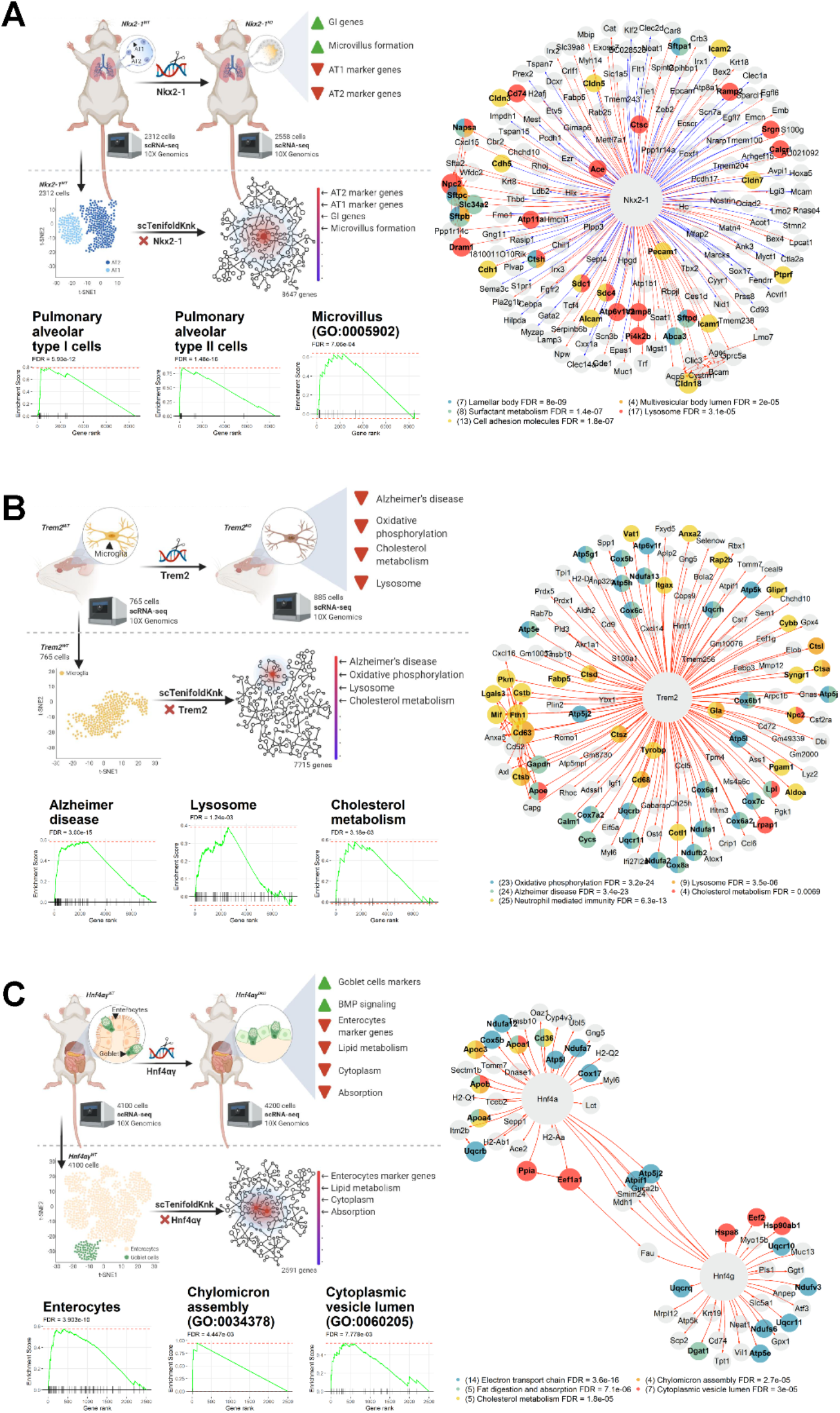
scTenifoldKnk virtual KO analysis recapitulates the findings of real KO experiments. (**A**) The schematic diagram (left top) shows the original KO experimental procedure and the virtual KO of *Nkx2-1*. GSEA plots (left bottom) show three enriched functions associated with virtual-KO perturbed genes upon the deletion of *Nkx2-1*. Gene rank indicates the position of each gene in the ranked gene list produced by scTenifoldKnk. The egocentric plot (right) shows the connections between the KO gene (*Nkx2-1*) and significant virtual-KO perturbed genes (FDR < 0.05). Nodes are color-coded by each gene’s membership association with enriched functional groups, as reported in the Enrichr analysis. The displayed gene sets were subsequently selected—i.e., only those with functions related to phenotypes reported in the corresponding original study are shown. The number of genes in the functional groups is given in the parentheses of the egocentric plot legend. (**B**) Same as **A** but for *Trem2*. (**C**) Same as **A** but for *Hnf4a* and *Hnf4g*.

#### Virtual vs. real KO experiment 2: Trem2 regulates microglial cholesterol metabolism

The Triggering Receptor Expressed on Myeloid cells 2 (*Trem2*) is a single-pass transmembrane immune receptor selectively expressed in microglia within the central nervous system. *Trem2* is known to be involved in late-onset Alzheimer’s disease and plays a role in modulating proliferation, survival, immune response, calcium mobilization, cytoskeletal dynamics, mTOR signaling, autophagy, and energy metabolism.^16^ The function of *Trem2* is known to be mediated via signaling transducer *Hcst* and adaptor *Tyrobp*.^17^ *Trem2* is also known to play a role in regulating lipid metabolism, with most studies focusing on lipids in the form of either lipoprotein particles or cell surface-exposed signals, such as candidate *Trem2* ligands.^18^ By comparing WT and *Trem2*^-/-^ KO mice, Poliani et al.^19^ showed that *Trem2* regulates many genes, such as*Apoe* and *Lpl*, which control lipid transport and catabolism in microglia. *Trem2* was also found to modulate gene expression of macrophages in adipose and control blood cholesterol metabolism in obese mice,^20^ further linking the function of *Trem2* to lipid metabolism. To examine whether *Trem2* mediates myelin lipid processing in microglia, Nugent et al.^21^ isolated and characterized *Cd11b*^+^/*Cd45^low^* microglial cells from *Trem2*^+/+^, *Trem2*^+/-^, and *Trem2*^-/-^ mice, fed with a 0.2% demyelinating cuprizone diet for 12 weeks. They analyzed a comprehensive set of analytical data using FACS, bulk RNAseq, scRNAseq, and lipidomics. They reported that *Trem2* upregulates *Apoe* and other genes involved in cholesterol transport and metabolism, causing robust intracellular accumulation of a storage form of cholesterol upon chronic phagocytic challenge. *Trem2* was also shown to regulate the expression of genes associated with cell damage response, lysosome and phagosome function, Alzheimer’s disease, and oxidative phosphorylation.^21^

To perform the virtual KO analysis, we obtained scRNAseq data from WT (*Trem2*^+/+^) mice.^21^ The expression matrix contained data of 7,715 genes and 765 *Cd11b*^+^/*Cd45^low^* microglial cells. We used this WT expression matrix as the input and used scTenifoldKnk to knock out *Trem2*. The final results of scTenifoldKnk analysis contained 128 virtual-KO perturbed genes (FDR < 0.05, **Supplementary Table S3**). The Enrichr analysis showed that the identified gene list was enriched with genes associated with *Alzheimer’s disease*, *oxidative phosphorylation*, *lysosome*, *TYROBP causal network*, *metabolic pathway of LDL*, *HDL and TG*, and *microglia pathogen phagocytosis pathway*. Such an enrichment indicates that the proteins attend to be functionally connected. These virtual-KO perturbed genes were highly interactive with each other, as shown by their positions on the STRING interaction network. The network of virtual-KO perturbed genes had significantly more interactions than expected (p-value < 0.01, STRING interaction enrichment test), which indicates that gene products exhibited more interactions among themselves than what would be expected for a random set of proteins of similar size, drawn from the genome. This result suggests that the identified virtual-KO perturbed genes are closely related to shared functions. Collectively, these scTenifoldKnk findings provide insight into understanding *Trem2* functions by revealing the list of genes perturbed following *Trem2* deletion (**Figure 3B**). These inferred functions are consistent with those reported in the original study.^21^

#### Virtual vs. real KO experiment 3: Hnf4a and Hnf4g stabilize enterocyte identity

Hepatocyte nuclear factor 4 alpha and gamma, *Hnf4a* and *Hnf4g*, are TFs that regulate gene expression in the gut epithelium. *Hnf4a* and *Hnf4g* function redundantly, and thus, an independent deletion of one paralog causes no gross abnormalities.^22,23^ *Hnf4a* and *Hnf4g* double-KO *Hnf4ag^DKO^* mice exhibit fluid-filled intestines indicative of an intestinal malfunction.^22^ Epithelial cells in the *Hnf4ag^DKO^* mutants fail to differentiate. Using bulk RNAseq data, Chen et al.^22^ compared gene expression in duodenal epithelial cells isolated from WT mice and *Hnf4a^KO^*, *Hnf4g^KO^* and *Hnf4ag^DKO^* mutants. They identified 2,892 DE genes in the *Hnf4ag^DKO^* mutant but only 560 and 77 in the *Hnf4a^KO^* and *Hnf4g^KO^* mutants, respectively [FDR < 0.05, absolute log2(fold change) > 1]. The DE genes identified in the *Hnf4ag^DKO^* enterocytes were enriched for functions in digestive metabolisms such as lipid metabolism, microvillus, and absorption, as well as enterocyte morphology, cytoplasm, Golgi apparatus, and immune signaling. *Hnf4ag^DKO^* epithelium exhibited a robust shift in the transcriptome away from differentiated cells toward proliferating and Goblet cells, suggesting that *Hnf4ag^DKO^* impairs enterocyte differentiation and destabilize enterocyte identity. To validate their findings, Chen et al.^22^ used scRNAseq to measure gene expression in intestinal villus epithelial cells. They obtained scRNAseq data for 4,100 and 4,200 cells from *Hnf4ag^WT^* and *Hnf4ag^DKO^* respectively and confirmed that compared to the WT, mutant epithelial cells show increased Goblet cell-enriched genes, such as *Agr2*, *Spink4*, *Gcnt3* and *S100a6*, and decreased expressions of enterocyte-enriched genes, such as *Npc1l1*, *Apoc3*, *Slc6a19* and *Lct* (**Figure 3C** left panel). The *Hnf4ag^DKO^* mutant cells also showed increased expression for genes in the BMP/SMAD signaling pathway and decreased expression of genes involved in lipid metabolism, microvillus and absorption, and genes related to the cytoplasm.^22^

We obtained the scRNAseq expression matrix of 4,100 cells from the *Hnf4ag^WT^* samples used as input for scTenifoldKnk. We constructed the WT scGRN of 2,591 genes and then virtually knocked out both *Hnf4a* and *Hnf4g* genes at the same time. The final result of scTenifoldKnk analysis contained 65 virtual-KO perturbed genes (FDR < 0.05, **Supplementary Table S4**). These genes were enriched with *electron transport chain, fat digestion and absorption*, *cholesterol metabolism*, *chylomicron assembly*, and *cytoplasmic vesicle lumen*. A search of the STRING database indicated that all virtual-KO perturbed genes form a fully connected network module. The interaction enrichment test showed that such a complete interconnection of 65 genes is less likely to be expected by chance (p-value < 0.01). Furthermore, GSEA analysis revealed that these virtual-KO perturbed genes were enriched with canonical marker genes of enterocytes (12 out of 132, **Figure 3C**), which is consistent with the finding of the original study.^22^

Interestingly, the number of significantly perturbed genes following the double-KO of *Hnf4a* and *Hnf4g*(65) was less than the single-KO of *Nkx2-1* (171) or *Trem2* (128). To examine the cause of this difference, we analyzed the relationship between the number of significantly perturbed genes and the degree (i.e., the number of connections) of the KO gene in the network (**Supplementary Figure S1**). We found a positive correlation between the two metrics (Pearson’s r=0.75, P = 0.03). *Hnf4a* and *Hnf4g* exhibited a lower degree (i.e., fewer connections) than *Nkx2-1* and *Trem2*, which may explain why KO of *Hnf4a* and *Hnf4g* produced fewer perturbed genes.

### Result 4: scTenifoldKnk virtual KO analysis detects functions of genes causative of Mendelian disorders

Mendelian diseases are a family of diseases caused by the loss or malfunctioning of a single gene. For many Mendelian diseases, we know a great deal about their genetic basis and pathophysiological phenotypes.^25^ We decided to use three Mendelian diseases, namely cystic fibrosis, Duchenne muscular dystrophy, and Rett syndrome, as “positive controls.” We tested the performance of scTenifoldKnk by determining whether it accurately predicted gene functions and hence inferred the molecular phenotypic consequences when the causative gene of each of these Mendelian diseases was virtually knocked out. As described in more detail below, in each case, we performed scTenifoldKnk analysis using existing scRNAseq data generated from cell types that are most relevant to the disease conditions (**Supplementary Table S1**).

#### Cystic fibrosis

Cystic fibrosis (CF) is one of the most common autosomal recessive diseases.^26^ It is caused by mutations in *CFTR*, a gene encoding for a transmembrane conductance regulator,^27,28^ which functions as a channel across the membrane of cells that produce mucus, sweat, saliva, tears, and digestive enzymes. The CFTR protein also regulates the function of other channels. *CFTR* is expressed in epithelial cells of many organs, including the lung, liver, pancreas, and digestive tract.^29^ The most common *CFTR* mutation that causes CF is the deletion of phenylalanine 508 (ΔF508), which disrupts the function of the chloride channels, preventing patients from regulating the flow of chloride ions and water across cell membranes.^27^ The truncated CFTR protein leads to a reduction of surfactants. It causes the build-up of sticky, thick mucus that clogs the airways, increasing the risk of bacterial infections and permanent lung damage.^30^

To test scTenifoldKnk, we obtained scRNAseq data from 7,326 pulmonary alveolar type II (AT2) cells in the GEO database (access number: GSM3560282). The original data sets were generated by Frank et al.^31^ to study the lineage-specific development of alveolar epithelial cells in mice. The original study was not directly focused on CF. Nevertheless, with the downloaded data, we constructed a WT scGRN that contained 7,107 genes. We then used scTenifoldKnk to knock out *Cftr*. The final results of scTenifoldKnk contained 17 virtual-KO perturbed genes: *Cftr*, *Birc5*, *Cldn10*, *Cxcl15*, *Dcxr*, *Hmgb2*, *Lamp3*, *Mgst1*, *Npc2*, *Pclaf*, *Pglyrp1*, *Sftpa1*, *Sftpb*, *Sftpc*, *Smc2*, *Tspan1* and *Tubb5* (FDR < 0.05, **Figure 4A, Supplementary Table S5**). Among them, *Sftpa1, Sftpb*, and *Sftpc* have functions associated with *ABC transporter disorders* and *surfactant metabolism. Sftpb* and *Sftpd* encode for surfactant proteins implicated in CF and innate immunity.^32,33^ *Ctsh* encodes for cathepsin H, a cysteine peptidase, which is involved in the processing and secretion of the pulmonary surfactant protein B.^34^ GSEA analysis of the ranked gene list was conducted using gene sets of the MGI mammalian phenotypes database as a reference. The result showed that the gene list scTenifoldKnk produced was significant in terms of *ion transmembrane transporter activity, abnormal surfactant secretion*, and *alveolus morphology* (FDR < 0.01 in all cases, **Figure 4A**). These results are consistent with the known pathophysiological changes resulting from the loss of *Cftr* function in the lungs.

**Figure 4.**
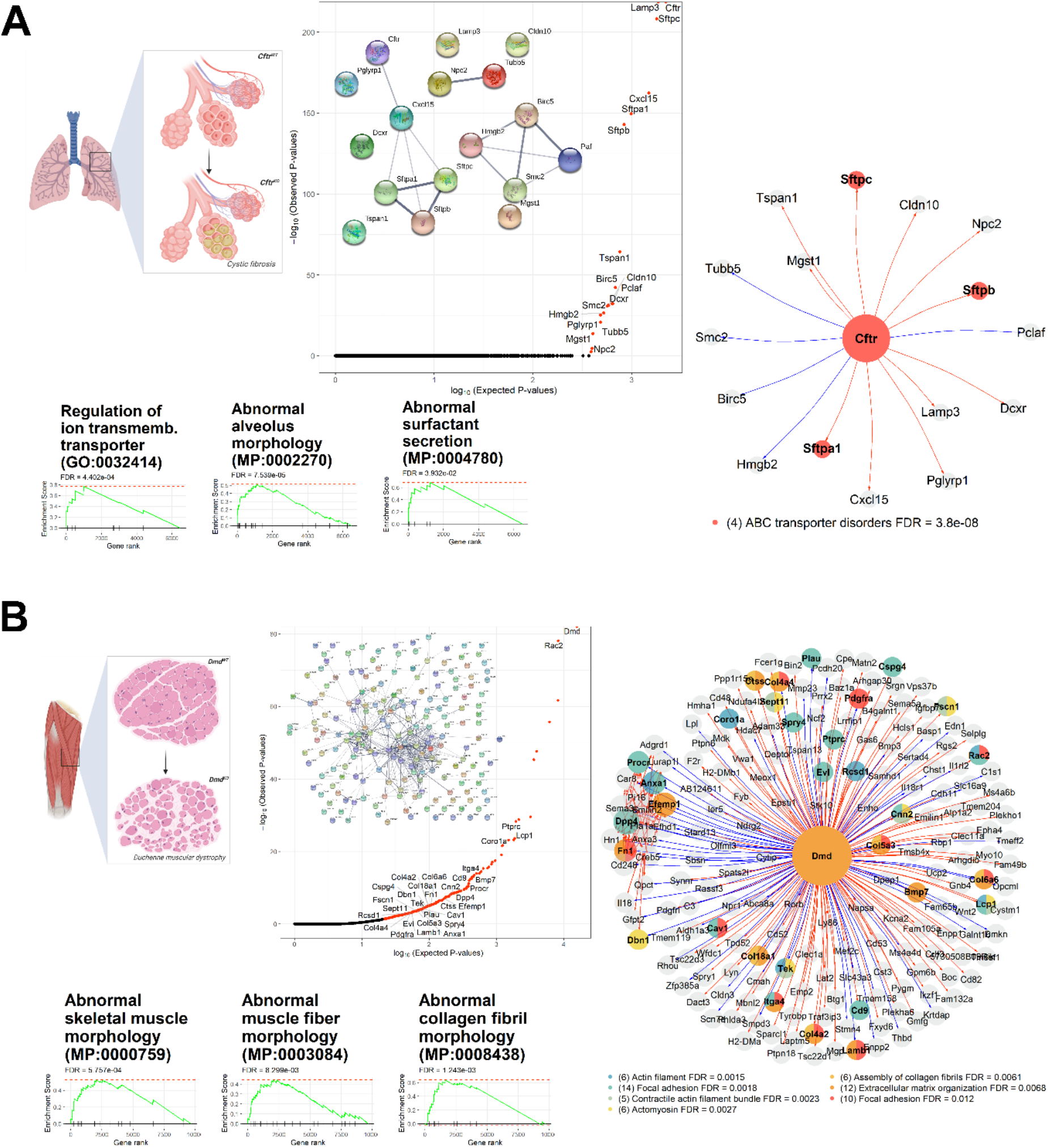
scTenifoldKnk virtual KO reveals functions of Mendelian disease genes in relevant cell types. (**A**) Virtual KO of *Cftr* in pulmonary alveolar type II cells identifies gene expression program changes associated with cystic fibrosis. GSEA analysis identifies significant gene sets, including *regulation of ion transmembrane transporter*, *abnormal alveolus morphology*, and *abnormal surfactant secretion*. Gene rank indicates the position of each gene in the ranked gene list produced by scTenifoldKnk. QQ-plot of genes and interconnection of virtual-KO perturbed genes in STRING are given. The egocentric plot (right) shows the connections between the KO gene (*Nkx2-1*) and significant virtual-KO perturbed genes (FDR < 0.05). Nodes are color-coded by each gene’s membership association with enriched functional groups, as reported in the Enrichr analysis. The displaying gene sets were subsequently selected—i.e., only those with functions related to the Mendelian disease phenotype are shown. (**B**) Virtual KO of *Dmd* in muscular cells identifies gene expression program changes associated with Duchenne muscular dystrophy. GSEA analysis identifies significant gene sets, including *abnormal skeletal muscle morphology* and *abnormal collagen fibril morphology*. Gene rank indicates the position of each gene in the ranked gene list produced by scTenifoldKnk. QQ-plot of genes and interconnection of virtual-KO perturbed genes in STRING are given. The egocentric plot (right) shows the connections between the KO gene (*Dmd*) and significant virtual-KO perturbed genes (FDR < 0.05). Nodes are color-coded by each gene’s membership association with enriched functional groups, as reported in the Enrichr analysis. The displaying gene sets are selected—only those with functions related to the Mendelian disease phenotype are shown.

To test the specificity of scTenifoldKnk, we decided to use genes that show highly similar expression profiles to that of *Cftr* to repeat the analysis. The selection of these genes was made by computing for each gene both the mean and variance of their expression levels across cells. Five closest genes to *Cftr—Akap7*, *Zranb1*, *Krcc1*, *Mta1* and *Rps8*—were selected. These selected genes may not necessarily have any functions related to that of *Cftr* in alveolar type II cells. When scTenifoldKnk was applied to predict perturbation profiles caused by their deletion, we found none of the sets of predicted functions similar to those specific functions (i.e., *ion transmembrane transporter function, abnormal surfactant secretion*, and *alveolus morphology*) associated with *Cftr*. Principal component analysis (PCA) confirmed that perturbation profiles of *Mta1*, *Akap7* and *Zranb1* differ most from that of *Cftr*, while the perturbation profiles of *Krcc1* and *Rps8* are relatively similar to that of *Cftr* (**Supplementary Figure S2**). Both *Krcc1* and *Rps8* have been reported to play a role in regulating cellular plasticity and fibrotic process.^35,36^

#### Duchenne muscular dystrophy

Duchenne muscular dystrophy (DMD) arises as a result of mutations in the open reading frame of *DMD*.^37,38^ The *DMD* gene encodes dystrophin, a large cytoskeletal structural protein, mostly absent in DMD patients.^39^ The absence of dystrophin results in a disturbance of the linkage between the cytoskeleton and the glycoproteins of the extracellular matrix, generating an impairment of muscle contraction, eventually leading to muscle cell necrosis (**Figure 4B**, left).^39,40^

We obtained the scRNAseq data of 5,159 muscle cells from the mouse limb (quadriceps) in the GEO database (GSM4116571). The original data was generated to study gene expression patterns in skeletal muscle cells.^41^ The original study was not focused on DMD. We used scRNAseq data from normal tissue to construct the WT scGRN of 9,783 genes. We subsequently performed the scTenifoldKnk virtual KO analysis to predict the molecular phenotype due to the impact of the *Dmd* KO. The final results of scTenifoldKnk included 190 virtual-KO perturbed genes (FDR < 0.05, **Supplementary Table S6**). These genes were enriched with functions related to *beta-1 integrin cell surface interaction*, *contractile actin filament bundle*, *actomyosin*, *extracellular matrix receptor interaction*, and *extracellular matrix organization* (**Figure 4B**, middle). GSEA analysis against the MGI mammalian phenotype database generated the following top hits (FDR < 0.01): *abnormal collagen fibril morphology*, *abnormal skeletal muscle morphology*, and *abnormal skeletal muscle fiber morphology* (**Figure 4B**, right). These phenotype terms are consistent with known effects of the loss of DMD function in muscle cells, verifying that scTenifoldKnk can predict phenotypic effects caused by gene KO pertinent to the biological context.

#### Rett syndrome

The third Mendelian disease we considered was the Rett syndrome (RTT, MIM 312750), which is a severe neurodevelopmental disease.^42,43^ RTT is known to be caused by mutations in *Mecp2*, a transcriptional repressor required to maintain normal neuronal functions.^44,45^ *Mecp2* deficiency in the brain decreases the expression level of genes involved in the *Bdnf* signaling pathway, mediated by repressing *Rest*.^46^

We obtained scRNAseq data generated from mouse neurons (SRA database access numbers: SRX3809326 and SRX3809327) for the mouse brain atlas project.^47^ The two data sets contain 2,054 and 2,156 neurons, respectively, derived from two CD1 P19 female mice that served as biological replicates. We analyzed the two data sets independently to see whether scTenifoldKnk could, as expected, produce similar results with data gathered from biological replicates. Two scGRNs containing 8,652 and 8,555 genes were constructed first. The scTenifoldKnk analysis of virtual KO of *Mecp2* produced 377 and 322 virtual-KO perturbed genes, respectively (FDR < 0.05, **Supplementary Table S7 & S8**), including 211 shared genes. The number of shared genes was significantly greater than random expectation (p-value < 10^-5^, hypergeometric test), indicating a high overlap rate between results of scTenifoldKnk analyses when applied to the two data sets from biological replicates. We also compared two ranked gene lists generated from the analysis of two replicate data sets. If scTenifoldKnk results are robust, we expected that the relative positions of the same genes in the two ranked lists should be more similar to each other than with a list of randomly ranked genes. Indeed, we found the correlation between rankings of two reported lists was positive (Spearman’s correlation coefficient ρ = 0.68) and highly significant (p-value < 10^-12^).

Many of these 211 genes were found to be targets of Rest. The enriched functions include axon, synaptic vesicle cycle, GABA synthesis, release, reuptake and degradation, syntaxin binding, and transmission across chemical synapses. GSEA analyses using ranked gene list as input showed BDNF signaling pathway was highly significant (FDR < 0.01, for both replicates, **Supplementary Figure S3**). These results are consistent with previous experimental results. For example, it is known that the most prominent alterations in gene products due to *Mecp2* KO are related to synapses and synaptic vesicle proteins.^48^ At the phenotypic level, the *Mecp2* KO causes early defects in GABAergic synapses and mediates autism-like stereotypies and RTT phenotypes.^49,50^ Mutations in syntaxin-1 are known to elicit phenotypes similar to those found in RTT.^51^

### Result 5: Additional evaluations of scTenifoldKnk-based virtual KO analysis

scTenifoldKnk attempts to infer the impact of knocking out a gene, given an input gene-by-cell expression matrix. This is done by first building a GRN and then “deleting” the gene from the GRN and aligning the resulting reduced GRN to the original GRN using manifold alignment. Manifold alignment is an essential step that cannot be replaced by simply picking up genes strongly linked with the KO gene. To confirm this, we revisited the outputs of *Trem2-KO* and *Dmd*-KO analyses to check whether DR genes are strongly linked with *Trem2* and *Dmd*, respectively. For *Trem2*, for example, we examined the correlation between each gene’s DR distance (estimated via manifold alignment after virtual KO of *Trem2*) and the edge weight of the link between the gene and *Trem2*. We found that the correlation was positive and significant (Spearman’s ρ = 0.69, p-value < 0.001), which was not unexpected as genes strongly linked with *Trem2* are supposed to be strongly impacted. However, the relationship is not linear—absolute values of the edge weights of most significant DR genes (i.e., genes with the largest DR distances) are variable, ranging from 0.075 to 0.40 (**Figure 5A**). The same trend was found with the *Dmd-KO* results (Spearman’s ρ = 0.53, p-value < 0.001, **Figure 5B**), with absolute values of the edge weights of most significant DR genes ranging from 0.025 to 0.37. These results indicate that significant DR genes identified using scTenifoldKnk are not necessarily always those strongly linked with the KO gene. Thus, scTenifoldKnk is not simply analyzing the adjacency matrix of the original GRN for the given KO gene to see which genes it is strongly connected to. Instead, many genes weakly linked with the KO were also identified as significant DR genes. We attributed this effect to the adoptaion of manifold alignment in scTenifoldKnk.

**Figure 5.**
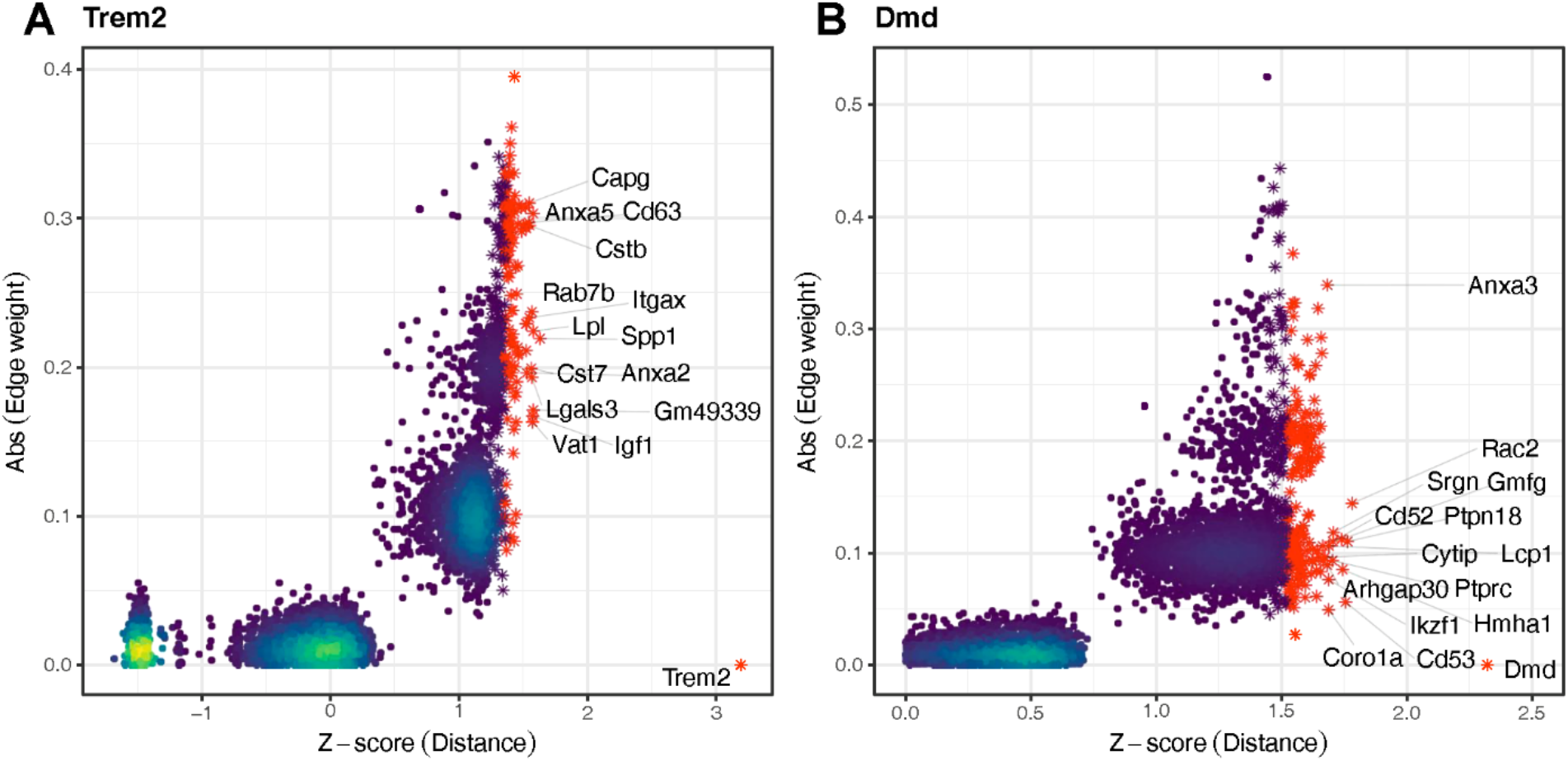
Correlation between genes’ DR distance and the edge weight of links between genes and the KO in scGRN. (**A**) Results from the microglial *Trem2-KO* analysis. Each dot represents a gene. The number of genes per unit area is shown with colors according to the density. The X-axis shows the Z score-normalized DR distance. For a given gene, the greater the distance, the more significant the gene is perturbed upon virtual KO. Significantly perturbed genes (FDR < 0.05) are shown in red asterisks. (**B**) Same as (**A**) but for *Dmd* in skeletal muscle cells.

Next, we note that scTenifoldKnk is designed to predict DR genes rather than differentially expressed (DE) genes. DR genes might be differentially expressed upon the gene KO. To examine the expression-level changes of virtual-KO perturbed genes, we performed a systematic comparison between the scTenifoldKnk results and the results of DE analysis across the three analyzed data sets: *Trem2*, *Nkx2-1*, and *Hnf4ag*. For each data set, we started by computing the DE statistics for all genes. Specifically, we obtained the fold change (FC) of each gene’s expression in WT samples related to KO samples (WT/KO) using the DE analysis package MAST.^24^ Then, we compared FC between significant DR genes and non-significant DR genes. We found that significant DR genes, or perturbed genes predicted by scTenifoldKnk, tend to have a larger FC value [p-value < 0.05 for all three cases (*Trem2*, *Nkx2-1*, and *Hnf4ag*), one-sided t-tests with log2-transformed FC values] than non-significant DR genes or non-perturbed genes (**Supplementary Figure S4**). Thus, the expression of DR genes predicted by scTenifoldKnk is more likely to be downregulated in samples of the real KO experiments.

Finally, we performed the analysis to evaluate the robustness of virtual KO analysis against cell sampling, using the microglial *Trem2* KO as an example. We randomly selected 500 cells each time and repeated the process ten times. For each subsampled data, we applied scTenifoldKnk to knock out *Trem2* and obtain a perturbation profile, i.e., a ranked list of genes sorted according to the DR distance. We found that the perturbation profiles of 10 subsamples are significantly positively correlated (average Spearman’s ρ = 0.55, **Supplementary Figure S5**). GSEA analyses with these perturbation profiles produced similar enriched gene sets.

### Result 6: Systematic KO of all genes individually in a given sample to obtain the KO perturbation profile landscape

As mentioned, scTenifoldKnk is designed and implemented to be computationally efficient. Indeed, we benchmarked the performance of scTenifoldKnk. After the WT scGRN is generated, scTenifoldKnk can complete virtual KO analysis at a rate of one minute per gene, as benchmarked on a PC with 2.9 GHz CPU and 16 GB RAM. That is to say, for a given data set of 6,000 genes, scTenifoldKnk can knock out all of these genes individually in 4 to 5 days. The process can also be split and run in parallel to increase the speed of computing. The outcome of such a systematic KO experiment is a collection of perturbation profiles of all genes. For each gene (e.g., gene *δ*), the perturbation profile of the gene (i.e., gene *δ*) is a vector of distances of all other genes ({*δ*^-^}) produced by scTenifoldKnk. The distance value quantifies the level of a gene (i.e., a gene in {*δ*^-^}) being perturbed by the deletion of the KO gene (i.e., gene *δ*). **Figure 6A** illustrates an analytic flowchart when scTenifoldKnk is used in a systematic KO experiment. For a given WT scGRN with *n* genes, scTenifoldKnk can be used to delete individual genes from 1 to n. For the *i*-th gene deletion, 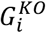, scTenifoldKnk produces a KO perturbation profile for the *i*-th gene. The KO perturbation profile is a vector of distances: [*d*_*i*,1_, *d*_*i*,2_,… *d_i,n_*]^*T*^, where *i* = 1, 2,…, n. Combining all genes’ KO perturbation profiles into an n × n matrix, called *KO perturbation profile matrix*, is followed by t-SNE embedding and clustering of genes.

**Figure 6.**
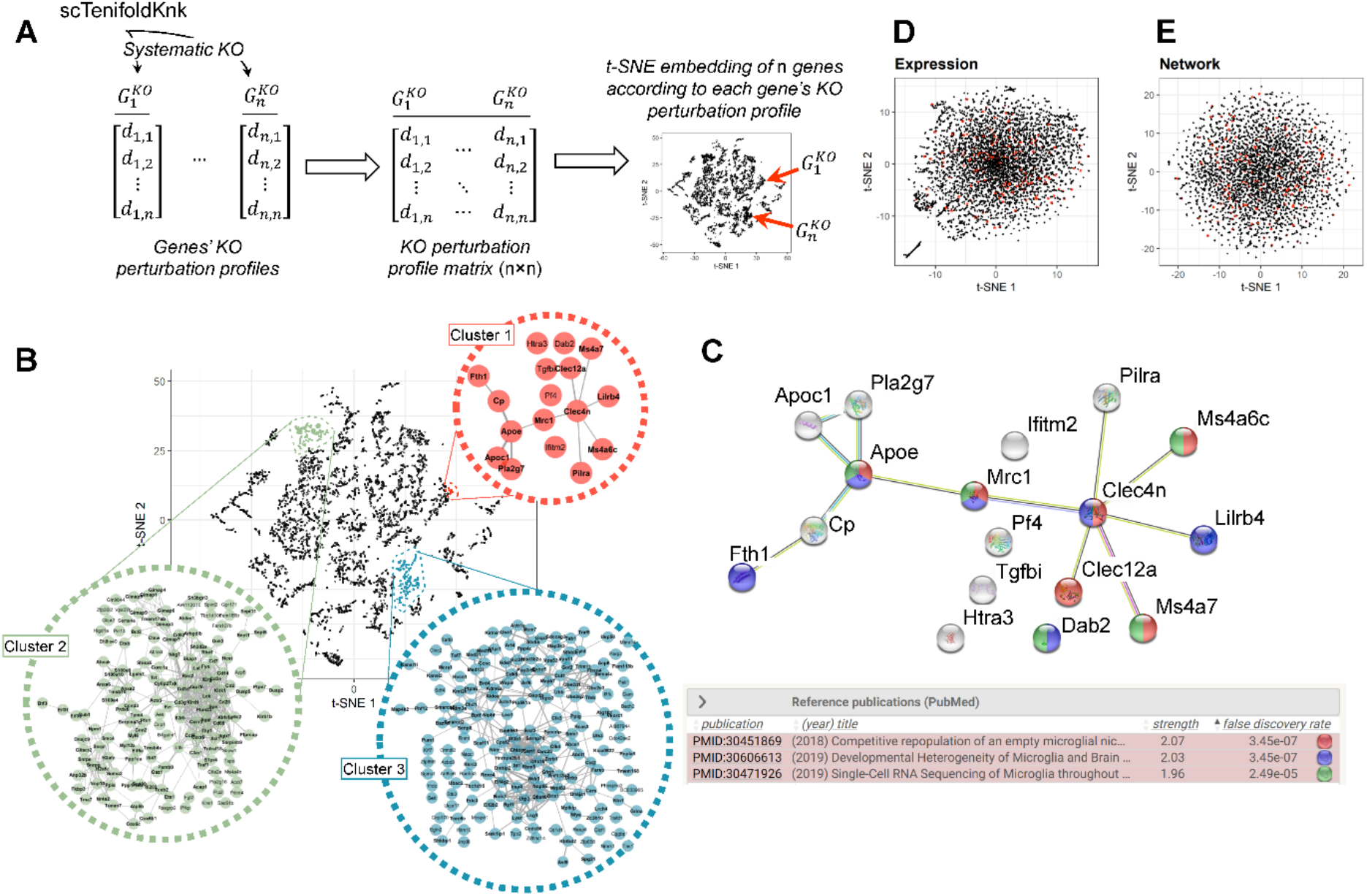
scTenifoldKnk enables a systematic KO experiment in microglia and the establishment of a KO perturbation profile landscape. (**A**) An illustration of systematic virtual-KO analysis using scTenifoldKnk. Two KO genes are shown as an example. Due to the difference in their profiles, two genes are embedded in two different locations, indicated by the red arrows, in the dimensionality reduction visualization. (**B**) t-SNE embedding of 6,853 genes expressed in microglia based on these genes’ perturbation profiles. Genes in three clusters in the embedding are highlighted. In the zoom-in view of each cluster, the connections between genes are retrieved from the STRING database. (**C**) STRING sub-network of 12 genes (*Apoc1*, *Apoe*, *Clec12a*, *Clec4n*, *Cp*, *Fth1*, *Lilrb4*, *Mrc1*, *Ms4a6c*, *Ms4a7*, *Pilra*, and *Pla2g7*) in Cluster 1 highlighted in **B**. References of three studies,^54–56^ in which the associations between genes were established, are given. (**D**) Same as **B** except the t-SNE embedding is based on genes’ expression profiles. The locations of genes in Cluster 1 are highlighted in red. (**E**) same as **B** except the t-SNE embedding is based on genes’ edge weight profile in the scGRN.

To demonstrate the use of the systematic KO functionality, we downloaded scRNAseq data from the brain immune atlas and obtained the expression matrix of 6,853 genes and 5,271 microglial cells (see **Experimental Procedures**). These microglial cells were derived from WT homeostatic mice.^52^ After knocking out all genes, we obtained the KO perturbation landscape of all 6,853 genes, as shown in the t-SNE embedding (**Figure 6B**). Each point represents a perturbation profile caused by the KO of a gene. Genes with similar perturbation profiles will be closely located in the low dimensional embedding. Therefore, examining genes situated closely will allow us to discover potential functional associations between them.^53^ We selected three clusters of different sizes to explore the member of genes in each cluster and the functional relationships between these genes (**Figure 6B**). We found that all three clusters contain genes that show a significantly higher level of functional associations than expected by chance (p-value < 0.01, STRING interaction enrichment tests). Cluster 1 contains 16 genes (**Figure 6C**).

According to the STRING database, 12 of these genes (*Apoc1*, *Apoe*, *Clec12a*, *Clec4n*, *Cp*, *Fth1*, *Lilrb4*, *Mrc1*, *Ms4a6c*, *Ms4a7*, *Pilra*, and *Pla2g7*) are functionally associated with each other, as supported by evidence from published literature.^54–56^ All these supporting references are related to the microglia study. The remaining four genes (*Htra3*, *Tgfbi*, *Pf4*, and *Ifitm2*) are not connected to this 12-gene sub-network.

Nevertheless, a new study showed that *Htra3* is overexpressed in repopulating microglia.^57^ Therefore, these “isolated” genes may be worthy of further scrutiny for their functions in microglia with the potential links to the other genes in the 12-gene sub-network. Cluster 2 contained 123 genes associated with the “Immune System Pathway” (FDR = 0.019, Reactome database), and Cluster 3 had 149 genes related to “Metabolism of RNA pathway” (FDR = 2e-4, Reactome database). Instead of using genes’ KO perturbation profiles to produce the landscape of the gene-gene relationship (as shown in **Figure 6B**), genes’ expression information can also be used to make such a landscape. Indeed, we applied t-SNE to the UMI count matrix and obtained the embedding plot of genes (**Figure 6D**). The difference between the embedding plot derived from the gene KO perturbation profile and the one derived from gene expression is noticeable (cf. **Figure 6B vs. 6D**). The latter had no structure among genes. Genes clustered together in the three example clusters in **Figure 6B** were found to be scattered in the embedding derived from gene expression (**Figure 6D**). Performing clustering on such an unstructured data cloud did not produce any meaningful results. Subsequently, we calculated the average distance between genes that belonged to the same KEGG gene sets. We found that, across all KEGG gene sets, the average distance in the embedding derived from KO perturbation profiles was significantly smaller than that in the embedding derived from the expression profile (**Supplementary Figure S6**). Similarly, an embedding plot of genes can be produced using genes’ network properties. We conducted additional analysis and generated the embedding derived from the genes’ profile of the weight of edges in the WT scGRN (**Figure 6E**). The same pattern was uncovered—that is, the average distance is smaller in the embedding derived from the KO perturbation profile than in that derived from the scGRN edge weight (**Supplementary Figure S6**). These results suggest that gene sets identified using the gene KO perturbation profile were more likely to be functionally connected than gene sets identified using other types of gene profiles.

### Result 7: Characterization of a multifunctional gene using scTenifoldKnk following systematic KO of the gene in multiple cell types

ScTenifoldKnk can be used to knock out a gene in different cell types systematically. In this way, it is possible to identify cell-type-specific functions of the gene and functions shared across multiple cell types. Here we use *MYDGF* (Myeloid-Derived Growth Factor) as an example KO gene to illustrate such an application. *MYDGF*, also known as *C19orf10*, is a 142-residue protein broadly expressed in multiple tissues and cell types.^58,59^ Mydgf has been shown in a mouse model to enhance cardiac myocyte survival, tissue repair, and angiogenesis caused by myocardial infarction.^60^ To elucidate *MYDGF*’s function, we downloaded multiple human scRNAseq data sets from the PanglaoDB database.^61,62^ From the downloaded data sets, we extracted cells from 45 different cell types. Subsequently, a virtual KO of *MYDGF* for each of these cell types and recovered 45 cell-type-specific perturbation profiles was performed. We first examined the perturbation profile of endothelial cells, in which the function of *MYDFG* has been studied.^63^ GSEA analysis with the ranked list of genes showed that the enriched functions of *MYDGF* include *cell cycle*, *VEGFA-VEGFR2 pathway*, and *Intra-Golgi traffic and activation* (**Figure 7A**). These results are consistent with previous findings,^63^ suggesting that scTenifoldKnk can recapitulate results from a cell-type-specific gene function study. We also concatenated perturbation profiles of 45 different cell types. There were 1,294 genes expressed across all tested cell types. From those genes, 364 were predicted as perturbed in all the cases. As expected, several cell types exhibited perturbation profiles similar to that of endothelial cells (**Figure 7B**). When using the Spearman’s ρ to measure the similarity between the perturbation profiles observed in endothelial cells and that observed in other cell types, these are the most similar cell types: T-cell (ρ = 0.59), hepatocyte (ρ = 0.45), podocyte (ρ = 0.33), and cardiomyocyte (ρ = 0.32). Indeed, previous studies show that Mydgf can regulate cell proliferation through the activation of Akt signaling pathways in those cells.^63–66^ In contrast, two cell types, proximal tubule cell (ρ = 0) and Leydig cell (ρ = −0.08), differed the most from endothelial cells, suggesting that Mydgf might play a very different role in these two cell types. In addition, a tSNE plot was produced to show the difference between cell types (**Supplementary Figure S7**). GSEA analysis identified enriched functions, including *AKT signaling*, *endothelial NOS activation*, *apoptosis*, *muscle contraction*, and *ALK2* pathway (**Figure 7C**).^67^ These findings are consistent with the fact that overexpression of *Mydgf* increases *AKT* phosphorylation and cell proliferation via *AKT/MAPK* signaling pathways.^66^ Identified functions were also related to *promoting survival and growth*, as previously reported.^60,68–70^ In summary, we used *Mydgf* as an example gene to demonstrate that scTenifoldKnk can be used to knock out a gene across multiple cell types in order to identify shared as well as cell type-specific functions of the KO gene.

**Figure 7.**
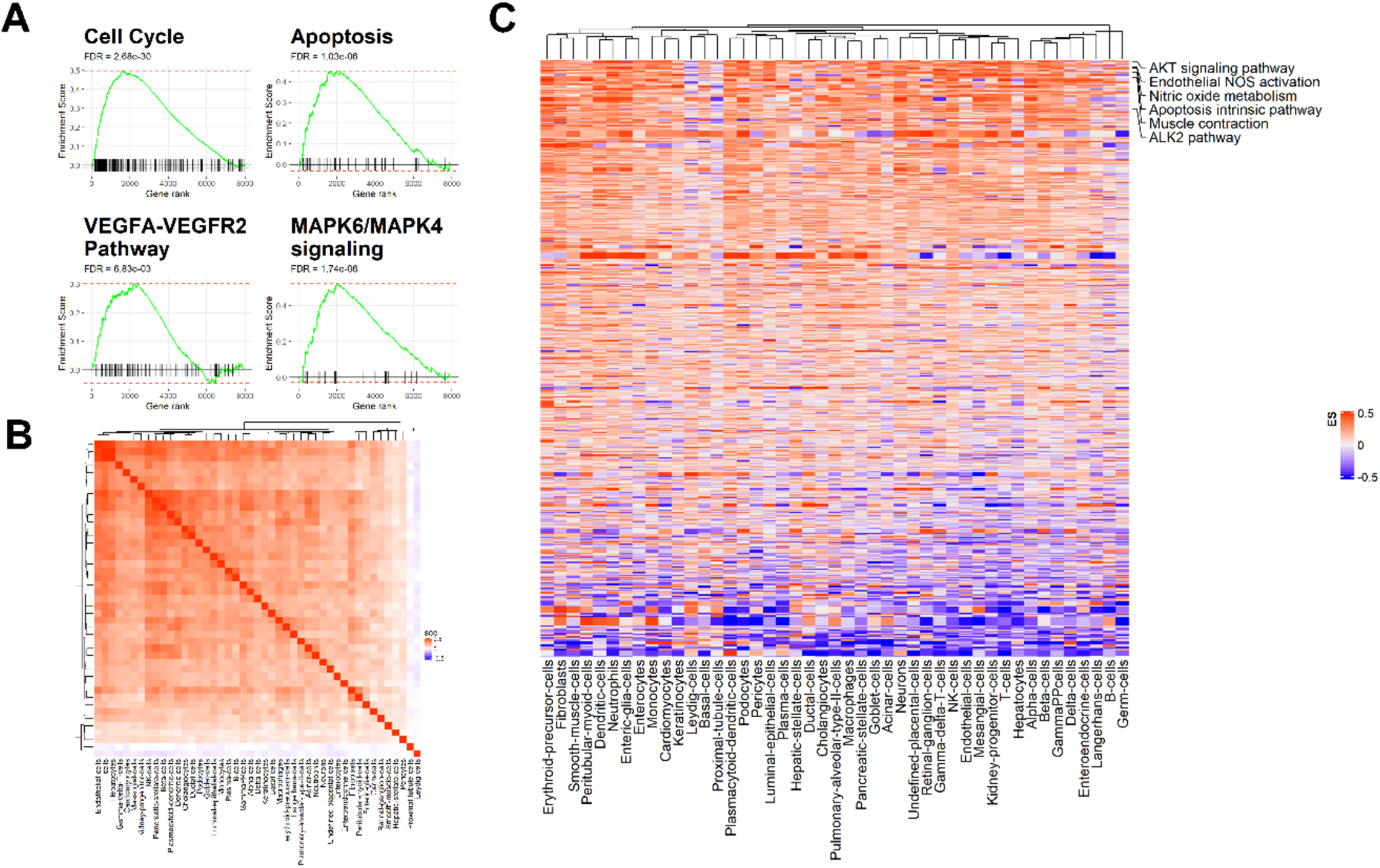
scTenifoldKnk virtual KO predicts shared and type-specific functions of *MYDGF* across cell types. (**A**) Results of GSEA analysis with ranked gene lists generated by virtual KO of *MYDGF* in endothelial cells. Significant gene sets include *cell cycle*, *apoptosis*, *VEGFA-VEGFR2 pathway*, and *MAPK6/4 signaling pathway*. (**B**) Correlation matrix showing similarity between *MYDGF* perturbation profiles of 45 cell types. (**C**) Heatmap showing GSEA enrichment scores for gene sets. Each row is a gene set in BioPlanet database. Rows are sorted in reverse order of average GSEA enrichment score across cell types. Names of several gene sets with functions known to be associated with *MYGDF* are shown.

## Discussion

Gene expression is almost always under coordinated regulation in cells of living organisms. Inferring GRNs is the key to a better understanding of such coordinated regulation. However, inferring GRNs is a challenging process—there are always many unknown variables in the system, and the power of inference is limited by the sample size. The development of single-cell technology has brought new “oil” to network science. We have previously shown that scRNAseq information can be leveraged to fuel the machine learning algorithms for reliable scGRN construction.^9^ In a GRN, the regulatory effect manifests as observable synchronized patterns of expression between genes. These genes are associated with the same biological process, pathway, or under the control of the same set of TFs.^71^ When a gene involved in a process is perturbed (e.g., knocked out), the expected first responders for such perturbation are those functionally closely related to the KO gene. Thus, modeling influence patterns in a GRN, such as using topological models to approximate perturbation patterns,^8^ can be used to predict gene function and prioritize target genes, which is helpful before expensive experimental measurements are undertaken. Thus, in principle, GRN-based perturbation analysis may contribute to the planning and designing of real-animal experimental work. Indeed, there is evidence showing that gene expression data need not necessarily be collected from perturbation experiments for GRN-based analysis to be successful.^72,73^

Our contribution is to provide a computationally efficient scGRN-based perturbation analytical system. By performing scTenifoldKnk virtual KO analyses with a series of existing scRNAseq data, we showed that scTenifoldKnk could predict gene function by identifying KO responsive genes. Overall, the inferred molecular functions of target genes are consistent with those enriched in genes reported in those original KO experiments. We also tested scTenifoldKnk using data from different cell types known to be affected in Mendelian diseases. While these diseases represent conditions caused by distinct genes and involve other dysregulated molecular processes, scTenifoldKnk demonstrated its value in all cases. Finally, we showed two case studies of systematic KO experiments.

Despite some apparent limitations associated with the virtual KO method, we start by discussing its advantages. First, the virtual KO method, as we implemented in scTenifoldKnk, is species agnostic—it works with scRNAseq data from humans and animal models. This feature gives the method a huge advantage for KO experiments focusing on human samples. In the lack of human KO samples, the KO animals are used as surrogates. The evolutionary divergence between humans and animal models is assumed to play a minor role in shaping orthologue gene function—but we know this is not always the case. While applying scTenifoldKnk to human scRNAseq data, researchers can avoid many pitfalls caused by extending the conclusions from animal KO experiments to humans. Second, scTenifoldKnk allows any gene to be knocked out for functional analysis as long as the gene expression is detectable in the WT sample. One may want to knock out all genes or a set of genes one by one to obtain a perturbation profile for each of the KO genes. Genes have similar perturbation profiles that are most likely to share molecular functions or are involved in the same signaling pathways. Genes with known functions can be used as positive controls to gauge the performance of scTenifoldKnk in the tested system. Third, scTenifoldKnk can be used to study the effects of gene KO across multiple cell types. Given that typical scRNAseq experiments generate expression data for various cell types, scTenifoldKnk can be used to predict the function of any KO gene in different cell types, allowing detection of diverse phenotypes associated with the KO gene. Finally, scTenifoldKnk can be used to study the function of essential genes, for which the gene KO causes lethal outcomes, making it impossible to establish the KO animals. When scTenifoldKnk is applied to these essential genes, especially with embryonic expression data of the genes from the WT samples, developmental functions of these genes can be studied.

scTenifoldKnk can be used in extended research areas beyond KO experiments. For example, biologists often need to know whether a genetic manipulation or perturbation will have an effect or not. scTenifoldKnk can be applied to make the prediction, suggest novel targets, and prioritize known targets before in vivo or in vitro studies. One may use drugs to block the transcription of predicted target genes in a candidate pathway. If the drug has an effect, one will conclude that the drug works on that pathway involved; otherwise, the pathway is not affected. The scTenifoldKnk-based analysis may apply to the follow-up research of genome-wide association studies (GWASs). GWASs have successfully detected associations between variants and phenotypes; however, a phenotypic trait is usually associated with many variants, presumably influencing gene expression regulation. scTenifoldKnk may be used to help geneticists to assess functional consequences to prioritize actionable gene targets. With example data sets, we showed that scTenifoldKnk could recapitulate major findings reported in real-KO experiments. We also showed the landscape of KO perturbation profiles of all genes in a given system. Experimentally, such systematic perturbation analysis has only been performed in yeast.^74^ With scTenifoldKnk, systematic KO analysis *in silico* can be performed in a cell-type-specific manner in any given organismal systems.

Limitations of scTenifoldKnk are inherited from being a virtual KO method. scTenifoldKnk cannot be used to predict the consequence of *gene overexpression*, which is also a commonly used method for gene function study. Also, as the power of scTenifoldKnk is rooted from the WT scGRN, the regulatory network from the WT sample, the prediction of scTenifoldKnk may “favor” regulatory rather than structural genes, as the latter tends to have smaller degree in the network. Nevertheless, it is still possible to adjust some details in the implementation of scTenifoldKnk to make it better fit user analytical needs. For example, instead of knocking out a target gene by setting its values to zeros in the adjacency matrix, a random shuffle of a gene’s expression values in the expression matrix may be used prior to the scGRN construction to mimic gene dysregulation.

Prediction of gene expression responses to perturbation using scRNAseq data is an active research area.^75^ To the best of our knowledge, there are two software tools that have been developed for this purpose: scGen and CellOracle.^6,7^ scGen is a package implemented in Python, using TensorFlow variational autoencoders combined with vector arithmetic to predict gene expression changes in cells.^6^ scGen works like a neural network-empowered regression tool that predicts the changes of gene expression in cells in response to specific perturbations such as disease and drug treatment. scGen requires training data sets from samples before and after being exposed to the same perturbation. CellOracle is a workflow, developed in Python with several R dependencies, that integrates scRNAseq and single-cell chromatin accessibility data (scATAC-seq) data to infer GRN and predict the changes of gene expression in response to specific perturbations. CellOracle constructs a GRN that accounts for the relationship between TFs and their target genes based on sequence motif analysis using the information provided by the scATAC-seq data. After that, the constructed GRN is further refined using regularized Bayesian regression models to remove weak connections and is adjusted to infer the context-dependent GRN using the scRNAseq data. Compared to scGen and CellOracle, scTenifoldKnk has a different, minimalistic design, specifically focusing on virtual KO. Unlike scGen, scTenifoldKnk does not need training data. Also, unlike CellOracle, scTenifoldKnk does not require information from scATAC-seq data.

Overall, we provide cogent evidence that scTenifoldKnk represents a powerful and efficient tool for conducting virtual KO analysis. The highly efficient implementation of scTenifoldKnk allows systematic deletion of many genes from any given scRNAseq data sets. The prediction power offered by scTenifoldKnk enables the accurate prediction of perturbations in regulatory networks caused by the deletion of a gene, so that the KO gene’s functions can be revealed in a cell type-specific manner. We anticipate that scTenifoldKnk will be adopted and widely applied in the predictions of gene function in single-cell biomedical research.

## Experimental Procedures

### Resource Availability

#### Lead Contact

Further information and requests for resources and reagents should be directed to and will be fulfilled by the lead contact, James J. Cai.

#### Materials Availability

This study did not generate new unique reagents.

#### Data and Code Availability

The source code of scTenifoldKnk is available at https://github.com/cailab-tamu/scTenifoldKnk. scTenifoldKnk has been implemented in R, Python, Julia, and Matlab. The R package is available at the CRAN repository at https://cran.r-project.org/web/packages/scTenifoldKnk/. The Matlab application is available in scGEAToolbox ^80^.

### The scTenifoldKnk workflow

scTenifoldKnk is a machine learning workflow for virtual KO experiments with scRNAseq data. It utilizes a scRNAseq expression matrix from a WT sample as input, without using any data from a KO sample, to predict regulatory network changes and perturbed genes caused by the KO of a gene. The input expression matrix is assumed to have been properly normalized.

### Construction of the WT scGRN

To construct scGRNs, scTenifoldKnk uses the method we previously proposed for scTenifoldNet.^9^ The procedure of scGRN construction consists of three steps: cell subsampling, principal component (PC) regression, and tensor decomposition.

#### (1) Cell subsampling

Initially, scTenifoldNet builds several subsets of cells via random sampling. Denote 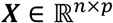 as the scRNAseq data matrix that reflects gene expression levels for *p* genes in *n* cells. A subsample set of cells is constructed via randomly sampling *m* (< *n*) cells in ***X***. By repeating this subsampling process for *t* times, *t* sub-sample sets of cells are derived, denoted as 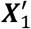,…, 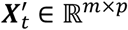.

#### (2) Network construction

For each 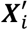, scTenifoldNet builds a GRN with an adjacency matrix ***W**^i^* via PC regression, where a PC analysis (PCA) is applied to the original explanatory variables, and then the response variable is regressed on a few leading PCs. Since PC regression only utilizes *d* PCs as the covariates in regression, where *d* << *min*(*m*, *n*), it mitigates over-fitting and reduces the computation time. To build an scGRN, each time scTenifoldNet focuses on one gene (referred to as the response gene) and applies PC regression. The expression level of the response gene is used as the response variable, and the expression levels of other genes are used as the explanatory variables in PC regression. scTenifoldNet repeats this process for another *p* - 1 times, with one different gene as the response gene each time. In the end, scTenifoldNet collects the coefficients of *p* regression models together and forms a *p* × *p* adjacency matrix ***W**^i^*, whose (*i*, *j*) entry saves the coefficient of the *i*-th gene on the *j*-th gene. ***W**^i^* could reflect the interaction strengths between each pair of genes.

#### (3) Network denoising

The adjacency matrices of the *t* networks ***W***^1^,…, ***W**^t^* can be stacked to form a third-order tensor 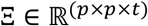. To remove noise and construct an overall adjacency matrix, scTenifoldNet applies CANDECOMP/PARAFAC (CP) tensor decomposition to Ξ to extract important latent factors. More specifically, scTenifoldNet approximates Ξ by Ξ^*R*^:

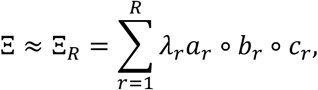

where ∘ denotes the outer product, 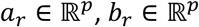, and 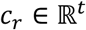 are unit-norm vectors, and *λ_r_* is a scalar. The reconstructed tensor 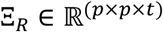 includes *t* denoised adjacency matrices, and by taking the average of them, scTenifoldNet obtains the overall stable adjacency matrix. After further normalizing its entries by dividing them by their maximum absolute value, scTenifoldNet generates the final adjacency matrix of scGRN for the given sample. For later use, denote it as ***W**_d_*.

#### (4) Adjusting edge weights of the directed network

We provided an option to adjust edge weights for the constructed scGRN. PC regression constructs directed networks. To obtain the strictly directed network, with a given, denoised network ***W**_d_*, for each gene pair (i, j) and (j, i), only the entry with a larger absolute weight is kept. More specifically, we defined the (i, j) entry for the strictly directed network ***W**_s_* by *W_s_*(i,j) = *W_d_*(i,j), if |*W_d_*(i,j)| > |*W_d_*(j,i)| and *W_s_*(i,j) = 0 otherwise. Note that if |*W_d_*(i,j)| < |*W_d_*(j,i)|, then we set *W_s_*(i,j) = 0, then the information in *W_d_*(i,j) is removed. To keep the information of *W_d_*(i,j), instead of removing the information completely, we defined a new parameter *λ*. Using this new parameter, we can integrate *W_d_* and *W_s_* to generate an “interpolated network” ***W**_i_*, which contains part of the information of *W_d_*(i, j). Given parameter *λ*, ***W**_i_* is define as:

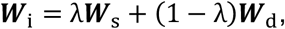

where *λ* ∈ [0,1]. It is easy to check that when *λ* = 0, we get the original denoised network ***W**_d_* and when *λ* = 1, it is to go back to the strictly directed network ***W**_s_*.

### Deletion of the KO gene from the WT scGRN

We propose the virtual KO method that directly works on the WT scGRN (**Figure 1B**). The adjacency matrix *W_d_* represents the scGRN constructed using the WT data. In the virtual KO method, the entire row of the adjacency matrix *W_d_* for the gene is set to zero. We denote the adjacency matrix of the scGRN generated as 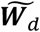.

### Comparison between the WT and pseudo-KO scGRNs

After obtaining ***W**_d_* and 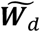, two comparable low-dimensional feature vectors of each gene in the two networks are built and then compared to detect affected genes. Our approach of creating low-dimensional feature vectors was inspired by manifold alignment and its application;^76–78^ our approach is referred to as quasi-manifold alignment because the adjacency matrices used here are not symmetric matrices while they are required to be symmetric in the original procedure. Here ***W**_d_* and 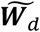 serve as the inputs for manifold alignment and the outputs are the low-dimensional features 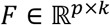 and 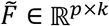 of genes before and after knocking out the target gene, where *k* << *p*. Before giving the details of the alignment procedure, we point out that ***W**_d_* and 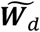 may include negative values, which reflect the negative correlation between genes. Before doing alignment, we add 1 to all entries in ***W**_d_* and 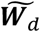, and the range of ***W**_d_* and 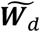 is transformed from [-1,1] to [0,2].

To perform quasi-manifold alignment, we first construct a joint adjacency matrix ***W*** by combining ***W**_d_* and 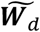 together, where 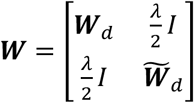. We can treat ***W*** as the adjacency matrix of a joint network formed by linking the corresponding genes in two networks. The off-diagonal block of this matrix reflects the corresponding genes between two networks. *λ* is a tuning parameter. In practice, we select *λ* as the mean of the row summations of ***W**_d_* and 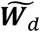. We further build 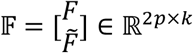, and the manifold alignment problem of two networks characterized by the adjacency matrices ***W**_d_* and 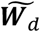 is equivalent to the manifold learning problem that finds the low dimensional features 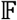 for the joint network characterized by the adjacency matrix ***W***. For the sake of convenience, we denote 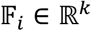 as the *i*-th row of 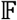 that reflects the projection corresponding to the *i*-th gene in the large network. The next step is to build a “Laplacian” matrix ***L*** = ***D*** – ***W***, where ***D*** is a diagonal matrix with 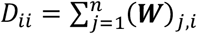. Denote *f*_1_, *f*_2_,…, *f_k_* as the eigenvectors corresponding to the *k* smallest non-zero eigenvalues of ***L***. Note that ***L*** is not a symmetric matrix. We found that the usual solution of symmetrizing ***L*** does not work well with either simulated or real data. We, therefore, use asymmetric matrix ***L*** in our quasi-manifold alignment procedure. Since ***L*** is not symmetric, there may be imaginary parts in the eigen decomposition. Based on our experiment, taking only the real part of eigenvectors with respect to the eigenvalue that has the smallest real part will give better overall results. The final low dimensional representation is 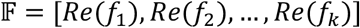, where *Re*(*f*) means the real part of *f*.

### Test for significance of virtual-KO perturbed genes

The virtual-KO perturbed genes are identified as genes with significant differences in their regulatory patterns in two scGRNs constructed from the WT and KO data. The method for testing the significance of the difference for each gene is described here. With 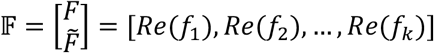 obtained in manifold alignment, for each gene, we calculate the distance *d_j_* between its two projected feature vectors from two networks. The rankings of *d_j_* are used to help identify significant genes. To avoid arbitrariness in deciding the number of selected genes, we proposed a χ^2^ test. Specifically, since 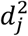 is calculated by taking the summation of squares of the differences of projected representations of two samples, its distribution could be approximately χ^2^. Instead of 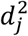, we use the scaled FC, 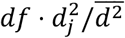, as the test statistic for each gene *j* to adjust the scale of the distribution, where *df* is the degree of freedom. 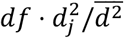 approximately follows a χ^2^-distribution with the *df* if the gene does not perform differently before and after knocking out the target gene. By using the upper tail (*P* [X>x]) of the χ^2^ distribution and the Benjamini–Hochberg (B–H) FDR correction for multiple testing correction,^79^ we assigned a p-value for each gene. To determine *df*, since the number of the selected significant genes will increase as *df* increases, we choose *df* = 1 to make a conservative selection of genes with high confidence.

### Gene functional annotation and enrichment tests

As described above, to predict the function of a given gene, the gene was virtually knocked out using scTenifoldKnk. The output of scTenifoldKnk represents the perturbation profile of the KO gene, which is a ranked list of virtual-KO perturbed genes, subject to Enrichr and GSEA functional enrichment tests.^10,14^ GSEA enrichment analysis was performed using the DR distances transformed using Box-Cox transformation and standardized using Z-score transformation. In brief, an enrichment score (ES) was calculated by walking down the list of genes ranked by the DR distance (proportional to the perturbation), increasing a running-sum statistic when we encounter a gene that is in a given gene set and decreasing when is not. The final enrichment score is equal to the maximum deviation from zero encountered in the random walk and corresponds to a weighted Kolmogorov-Smirnov-like statistic. The reference gene sets used for functional enrichment tests included *KEGG_2019_Human*, *KEGG_2019_Mouse*, *GO_Biological_Process_2018*, *GO_Cellular_Component_2018*, *GO_Molecular_Function_2018*, *BioPlanet_2019*, *WikiPathways_2019_Human*, *WikiPathways_2019_Mouse*, and *Reactome_2016*. Top genes in the ranked list were tested for significance (see above), and significant genes were used as input for the Enrichr analysis ^14^. The full ranked gene list was used as input for GSEA analysis ^10^. To remove redundant results produced by the GSEA analysis, identified significant functional terms and gene sets were grouped according to the overlap of leading-edge genes. More specifically, for a given pair of identified gene sets, if Jaccard index, 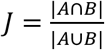, is greater than 0.8, where A and B are leading-edge genes of the two gene sets, then this pair of gene sets were grouped together. The functional annotation of grouped gene sets was reported under the same enriched function group. In this way, non-redundant function groups of identified gene sets were created and used for functional inference of the KO gene. GSEA analysis was also conducted against gene sets made of marker genes obtained from PanglaoDB.^61^ The protein interaction enrichment tests were performed using the web tool provided by the STRING database.^15^ The CSV files of gene set enrichment results are available for downloading from the Github website of scTenifoldKnk.

### Systematic KO analysis

Systematic KO analysis was performed with the microglial scRNAseq data from the brain immune atlas. Data used in the systematic KO analysis was obtained from the brain immune atlas (https://www.brainimmuneatlas.org), a scRNAseq resource for assessing and capturing the diversity of the brain immune compartment, as published in ref.^52^ Data was generated using the 10x Genomics Chromium platform, including more than 61,000 CD45+ immune cells from whole brains or isolated dura mater, subdural meninges, and choroid plexus of mice. The downloaded data, referred to as the *full aggregate data set* (combining cells of whole brain and choroid plexus cells from WT + Irf8 KO mice), was stored in the file named filtered_gene_bc_matrices_mex_irf8_fullAggr.zip. The downloaded matrix was processed, and the sub-matrix contained 5,271 microglia from the WT mice. For all genes, the KO perturbation profile of each gene, i.e., a vector of DR distances, was transformed using Box-Cox transformation and then was standardized using z-score transformation. The processed KO perturbation profiles of all genes were combined into one matrix for t-SNE embedding.

## Acknowledgments

A sincere thank you to Dr. Jingshu Chen and Dr. Andrew Hillhouse for technical support and the editing team members of the English 320 course project, Kamryn Watson, Kanza Akhtar, Marisa Morris, and Mary Frances Nance, for proofreading of this paper. This research was funded by Texas A&M University X-Grant (2019), T3-Grant (2019), and DoD GW200026 for J.J.C., the Allen Endowed Chair in Nutrition & Chronic Disease Prevention, NIH R01-ES025713, R01-CA202697, R35-CA197707 grants for R.S.C., and Texas A&M Institute of Data Science (TAMIDS) Data Resource Development Program Award (2020) for D.O. and Y.Z.

## Author Contributions

J.J.C. conceived the study, designed the workflow, and implemented the Matlab version of the software. D.O. designed the workflow, performed data analysis, and implemented the R version of the software. Y.Z. and G.L. contributed to the workflow design. Q.X. and Y.Y. contributed to the preparation and analysis of testing data sets. Y.T., R.S.C., and J.Z.H contributed to the concept development and the writing of the manuscript. J.J.C. and J.Z.H. supervised the data analysis.

## Declaration of Interests

D.O., Y.Z., G.L., Y.Y, J.Z.H. and J.J.C. are listed as inventors on a patent application related to this work.

## Supplementary Tables

**Supplementary Table S1. Summary of real-data applications of scTenifoldKnk analysis.**

**Supplementary Table S2. 171 genes perturbed by the virtual-KO of Nkx2-1 in alveolar cells.** The STRING interaction network of perturbed genes is available at the permalink: https://version-11-0b.string-db.org/cgi/network?networkId=bECocSTVt8bZ

**Supplementary Table S3. 128 genes perturbed by the virtual-KO of Trem2 in microglial cells.** The STRING interaction network of perturbed genes is available at the permalink: https://version-11-0b.string-db.org/cgi/network?networkId=bEZKYpHcHsns

**Supplementary Table S4. 65 genes perturbed by the virtual-KO of Hnf4a and Hnf4g in intestinal cells.** The STRING interaction network of perturbed genes is available at the permalink: https://version-11-0b.string-db.org/cgi/network?networkId=bGGZHJwMOcny

**Supplementary Table S5. 17 genes perturbed by the virtual-KO of Cftr in alveolar type II cells.** The STRING interaction network of perturbed genes is available at the permalink: https://version-11-0b.string-db.org/cgi/network?networkId=bbh2WkHUgVGT

**Supplementary Table S6. 190 genes perturbed by the virtual-KO of Dmd in skeletal muscle cells of the mouse limb.** The STRING interaction network of perturbed genes is available at the permalink: https://version-11-0b.string-db.org/cgi/network?networkId=bY1kGZIAZ55e

**Supplementary Table S7. 377 genes perturbed by the virtual-KO of Mecp2 in mouse neurons (Biological replicate 1).** The STRING interaction network of perturbed genes is available at the permalink: https://version-11-0b.string-db.org/cgi/network?networkId=bUTs7s0SjR9E

**Supplementary Table S8. 322 genes perturbed by the virtual-KO of Mecp2 in mouse neurons (Biological replicate 2).** The STRING interaction network of perturbed genes is available at the permalink: https://version-11-0b.string-db.org/cgi/network?networkId=bRZ7FLRdM1Ic

## Notes

https://github.com/cailab-tamu/scTenifoldKnk

### Summary of Updates

We added a systematic validation of the predictions using differential expression analysis. New figures and tables are presented.

