## Supplementary Material for "scTenifoldKnk: an efficient virtual knockout tool for gene function predictions via single-cell gene regulatory network perturbation"

### Supplementary Data

Supplementary Table S4. 65 genes perturbed by the virtual-KO of *Hnf4a* and *Hnf4g* in intestinal cells. 8

#### Supplementary Tables

##### Supplementary Table S1. Summary of real-data applications of scTenifoldKnk analysis.

| | KO gene | Cell type | Main results from original and related studies | Original study | **scTenifoldKnk predicted results** (i.e., enriched functions of virtual-KO perturbed genes) | **Figure and table in this article** (Number of virtual-KO perturbed genes) |
|---|---|---|---|---|---|---|
| Real-animal KO experiments with published data | | | | | | |
| 1 | Nkx2-1 | Pulmonary alveolar cells | Decreased expression of marker genes of alveolar type I (AT1) & II (AT2) cells; Increased expression of marker genes of gastrointestinal cells; *Nkx2-1* regulates expression of genes related to membrane composition, extracellular matrix, and cytoskeleton; mutations in Nkx2-1 interrupts AT2 cell function and identity [1, 2] | [1] | Marker genes of AT1 and AT2 cells; intestinal microvillus, cell cycle, and cytoskeleton; epithelial to mesenchymal transition led by the WNT signaling pathway members; surfactant homeostasis; lamellar body; cell adhesion molecules | **Figure 3A**; **Suppl. Table S2** (171 genes) |
| 2 | Trem2 | Microglial cells | *Trem2* regulates expression of genes related to lipid transport and catabolism [3], lipid metabolism [4], microglial cell damage response [5], lysosome and phagosome function, Alzheimer's disease, and oxidative phosphorylation [6]; *Trem2* interacts with signaling transducer *Hcst* and adaptor *Tyrobp* [7] | [6] | Alzheimer's disease; oxidative phosphorylation; lysosome; TYROBP causal network; microglia pathogen phagocytosis pathway | **Figure 3B**; **Suppl. Table S3** (128 genes) |
| 3 | Hnf4a & Hnf4g | Intestinal villus epithelial cells | Increased expression of Goblet cell-enriched genes (e.g., *Agr2, Spink4, Gcnt3* and *S100a6*) and genes in BMP/SMAD signaling pathway; decreased expression of enterocyte-enriched genes (e.g., *Npc1l1, Apoc3, Slc6a19* and *Lct*) genes involved in lipid metabolism, microvillus and absorption, and genes related to cytoplasm [8] | [8] | Enterocyte marker genes; electron transport chain; fat digestion and absorption; cholesterol metabolism; chylomicron assembly; cytoplasmic vesicle lumen | **Figure 3C**; **Suppl. Table S4** (65 genes) |
| Mendelian diseases | | | | | | |
| 4 | *Cftr* (Cystic fibrosis) | Pulmonary alveolar cells | Expressed in epithelial cells of many organs [9]; mutations disrupt the function of the chloride channels, preventing them from regulating the flow of chloride ions and water across cell membranes [10, 11] | [12] | ABC transporter disorders and surfactant metabolism; ion transmembrane transporter activity, abnormal surfactant secretion, and alveolus morphology | **Figure 4A**; **Suppl. Table S5** (17 genes) |

| | | | | | | |
|---|---|---|---|---|---|---|
| 5 | *Dmd* (Duchenne muscular dystrophy) | Skeletal myocytes | Mutation disrupts the linkage between the cytoskeleton and the glycoproteins of the extracellular matrix; impairment of muscle contraction; muscle cell necrosis [13, 14] | [15] | beta-1 integrin cell surface interaction, contractile actin filament bundle, actomyosin, extracellular matrix receptor interaction, extracellular matrix organization; abnormal collagen fibril morphology, abnormal skeletal muscle morphology, abnormal skeletal muscle fiber morphology | **Figure 4B**; **Suppl. Table S6** (190 genes) |
| 6 | *Mecp2* (Rett syndrome) | Neurons | Decreases the expression level of the genes involved in the BDNF signaling [16]; repressing the TF REST [16]; dysregulated maintenance of normal neuronal functions [17, 18]; altered synapses and synaptic vesicle proteins [19]; defects of GABAergic synapses [20]; autism-like stereotypies [21]; syntaxin-1 mutant phenotype [22] | [23] | Affected *REST* target genes; synaptic vesicle cycle, GABA synthesis, release, reuptake and degradation, syntaxin binding, transmission across chemical synapses | **Suppl. Figure S1**; **Suppl. Table S7 and S8** (377 and 322 genes, respectively) |

#### Supplementary Table S3. 128 genes perturbed by the virtual-KO of *Trem2* in microglial cells.

The STRING interaction network of perturbed genes is available at the permalink: https://version-11-0b.string-db.org/cgi/network?networkId=bEZKYpHcHsns

| GENE | DISTANCE | Z | FC | P.VALUE | P.ADJ |
|---|---|---|---|---|---|
| *Trem2* | 0.000138 | 3.194156 | 5471989 | 0 | 0 |
| *Spp1* | 6.22E-07 | 1.631 | 110.8313 | 6.44E-26 | 2.45E-22 |
| *Lpl* | 5.02E-07 | 1.579041 | 72.12195 | 2.02E-17 | 4.17E-14 |
| *Gm49339* | 5.01E-07 | 1.578712 | 71.92444 | 2.24E-17 | 4.17E-14 |
| *Anxa5* | 5.00E-07 | 1.57804 | 71.52298 | 2.74E-17 | 4.17E-14 |
| *Igf1* | 4.92E-07 | 1.574502 | 69.44577 | 7.85E-17 | 9.95E-14 |
| *Vat1* | 4.88E-07 | 1.572462 | 68.27466 | 1.42E-16 | 1.54E-13 |
| *Rab7b* | 4.80E-07 | 1.568251 | 65.91764 | 4.70E-16 | 4.47E-13 |
| *Lgals3* | 4.79E-07 | 1.567807 | 65.67373 | 5.32E-16 | 4.49E-13 |
| *Cst7* | 4.70E-07 | 1.56323 | 63.21053 | 1.86E-15 | 1.41E-12 |
| *Cstb* | 4.44E-07 | 1.549802 | 56.48813 | 5.65E-14 | 3.91E-11 |
| *Itgax* | 4.41E-07 | 1.548187 | 55.72814 | 8.32E-14 | 5.27E-11 |
| *Capg* | 4.40E-07 | 1.547522 | 55.41804 | 9.74E-14 | 5.70E-11 |
| *Cd63* | 4.31E-07 | 1.54291 | 53.31282 | 2.84E-13 | 1.54E-10 |
| *Anxa2* | 4.06E-07 | 1.528458 | 47.2053 | 6.39E-12 | 3.24E-09 |
| *Aplp2* | 4.02E-07 | 1.526315 | 46.35953 | 9.84E-12 | 4.68E-09 |
| *Apoe* | 3.98E-07 | 1.52393 | 45.43527 | 1.58E-11 | 6.79E-09 |
| *Cxcl16* | 3.98E-07 | 1.52383 | 45.39689 | 1.61E-11 | 6.79E-09 |
| *Cd72* | 3.94E-07 | 1.521539 | 44.52685 | 2.51E-11 | 1.00E-08 |
| *Cd52* | 3.92E-07 | 1.520267 | 44.05086 | 3.20E-11 | 1.22E-08 |
| *Mif* | 3.63E-07 | 1.502295 | 37.8287 | 7.72E-10 | 2.80E-07 |
| *Tmsb10* | 3.42E-07 | 1.488064 | 33.5148 | 7.07E-09 | 2.44E-06 |
| *Pkm* | 3.35E-07 | 1.482903 | 32.07115 | 1.49E-08 | 4.91E-06 |
| *Ccl6* | 3.10E-07 | 1.464961 | 27.50665 | 1.57E-07 | 4.96E-05 |
| *Fth1* | 3.09E-07 | 1.464568 | 27.4142 | 1.64E-07 | 4.99E-05 |
| *Atp5g1* | 3.02E-07 | 1.459181 | 26.17544 | 3.12E-07 | 8.99E-05 |
| *Atp5k* | 3.02E-07 | 1.458958 | 26.12521 | 3.20E-07 | 8.99E-05 |
| *Axl* | 3.02E-07 | 1.458659 | 26.05818 | 3.31E-07 | 8.99E-05 |
| *Lyz2* | 3.01E-07 | 1.458107 | 25.935 | 3.53E-07 | 9.25E-05 |
| *Cybb* | 2.92E-07 | 1.4513 | 24.46022 | 7.59E-07 | 0.000192 |
| *Adssl1* | 2.91E-07 | 1.450387 | 24.26864 | 8.38E-07 | 0.000205 |
| *Crip1* | 2.84E-07 | 1.444739 | 23.11578 | 1.53E-06 | 0.000362 |
| *Tpm4* | 2.83E-07 | 1.444165 | 23.00159 | 1.62E-06 | 0.000373 |
| *Ch25h* | 2.82E-07 | 1.443426 | 22.85547 | 1.75E-06 | 0.00039 |
| *Ctsb* | 2.81E-07 | 1.442224 | 22.61972 | 1.97E-06 | 0.000429 |
| *Aldoa* | 2.73E-07 | 1.435682 | 21.37695 | 3.77E-06 | 0.000796 |
| *Pld3* | 2.72E-07 | 1.434852 | 21.22423 | 4.09E-06 | 0.000839 |
| *Glipr1* | 2.71E-07 | 1.434119 | 21.09015 | 4.38E-06 | 0.000876 |
| *Tyrobp* | 2.70E-07 | 1.433059 | 20.89753 | 4.85E-06 | 0.000944 |
| *Cd9* | 2.69E-07 | 1.432514 | 20.79925 | 5.10E-06 | 0.000969 |
| *Ndufa1* | 2.68E-07 | 1.431044 | 20.53646 | 5.85E-06 | 0.001085 |
| *Ccl5* | 2.66E-07 | 1.429248 | 20.21954 | 6.90E-06 | 0.001249 |
| *Fabp5* | 2.65E-07 | 1.428532 | 20.09453 | 7.37E-06 | 0.001303 |
| *Ass1* | 2.59E-07 | 1.423386 | 19.21793 | 1.17E-05 | 0.002014 |

| | | | | | |
|---|---|---|---|---|---|
| *Cox6a2* | 2.56E-07 | 1.420911 | 18.8098 | 1.44E-05 | 0.002426 |
| *Prdx1* | 2.56E-07 | 1.420717 | 18.77815 | 1.47E-05 | 0.002426 |
| *Fabp3* | 2.55E-07 | 1.420203 | 18.69456 | 1.53E-05 | 0.002458 |
| *S100a1* | 2.55E-07 | 1.419846 | 18.63672 | 1.58E-05 | 0.002458 |
| *Cox6a1* | 2.55E-07 | 1.419821 | 18.6326 | 1.58E-05 | 0.002458 |
| *Cops9* | 2.51E-07 | 1.416091 | 18.03869 | 2.16E-05 | 0.00329 |
| *Plin2* | 2.49E-07 | 1.414001 | 17.71387 | 2.57E-05 | 0.003826 |
| *Rhoc* | 2.48E-07 | 1.413359 | 17.61525 | 2.70E-05 | 0.003952 |
| *Tceal9* | 2.48E-07 | 1.413063 | 17.56992 | 2.77E-05 | 0.003971 |
| *Gapdh* | 2.45E-07 | 1.4103 | 17.15258 | 3.45E-05 | 0.004855 |
| *Tmem256* | 2.44E-07 | 1.409245 | 16.99577 | 3.75E-05 | 0.005177 |
| *Atp5l* | 2.43E-07 | 1.408756 | 16.92361 | 3.89E-05 | 0.005281 |
| *Tomm7* | 2.42E-07 | 1.407873 | 16.79394 | 4.17E-05 | 0.005555 |
| *Cd68* | 2.41E-07 | 1.406806 | 16.63853 | 4.52E-05 | 0.005926 |
| *Gnas* | 2.40E-07 | 1.405821 | 16.49648 | 4.87E-05 | 0.00621 |
| *Fxyd5* | 2.40E-07 | 1.405745 | 16.48546 | 4.90E-05 | 0.00621 |
| *Bola2* | 2.39E-07 | 1.40497 | 16.37448 | 5.20E-05 | 0.006476 |
| *Cxcl14* | 2.37E-07 | 1.402903 | 16.08204 | 6.07E-05 | 0.007343 |
| *Pgk1* | 2.37E-07 | 1.402856 | 16.07551 | 6.09E-05 | 0.007343 |
| *Aldh2* | 2.36E-07 | 1.40155 | 15.89341 | 6.70E-05 | 0.007958 |
| *Prdx5* | 2.33E-07 | 1.399249 | 15.57743 | 7.92E-05 | 0.00926 |
| *Cox6c* | 2.33E-07 | 1.399007 | 15.54457 | 8.06E-05 | 0.009279 |
| *Ifitm3* | 2.32E-07 | 1.398301 | 15.44901 | 8.48E-05 | 0.009615 |
| *Atp5j2* | 2.31E-07 | 1.396981 | 15.27187 | 9.31E-05 | 0.010301 |
| *Elob* | 2.31E-07 | 1.396915 | 15.26309 | 9.35E-05 | 0.010301 |
| *Atp5mpl* | 2.27E-07 | 1.392628 | 14.70181 | 0.000126 | 0.013624 |
| *Sem1* | 2.26E-07 | 1.392334 | 14.66407 | 0.000128 | 0.013624 |
| *Uqcr11* | 2.26E-07 | 1.392266 | 14.65529 | 0.000129 | 0.013624 |
| *Ifi27l2a* | 2.26E-07 | 1.391984 | 14.61927 | 0.000132 | 0.013697 |
| *Ctsd* | 2.26E-07 | 1.391717 | 14.58514 | 0.000134 | 0.013759 |
| *Calm1* | 2.25E-07 | 1.391513 | 14.55911 | 0.000136 | 0.013764 |
| *Atox1* | 2.24E-07 | 1.389937 | 14.35975 | 0.000151 | 0.0151 |
| *Cox5b* | 2.23E-07 | 1.389507 | 14.30581 | 0.000155 | 0.015337 |
| *Ndufb2* | 2.23E-07 | 1.389299 | 14.2797 | 0.000158 | 0.015351 |
| *Cox7a2* | 2.22E-07 | 1.388302 | 14.15565 | 0.000168 | 0.016073 |
| *Cox7c* | 2.22E-07 | 1.38801 | 14.11951 | 0.000172 | 0.016073 |
| *Atp5e* | 2.22E-07 | 1.387978 | 14.11548 | 0.000172 | 0.016073 |
| *Cox6b1* | 2.22E-07 | 1.387845 | 14.09914 | 0.000173 | 0.016073 |
| *Myl6* | 2.22E-07 | 1.387654 | 14.07561 | 0.000176 | 0.01608 |
| *Arpc1b* | 2.21E-07 | 1.387222 | 14.02235 | 0.000181 | 0.016345 |
| *Ybx1* | 2.21E-07 | 1.386869 | 13.97906 | 0.000185 | 0.016406 |
| *Ndufa2* | 2.21E-07 | 1.386804 | 13.97111 | 0.000186 | 0.016406 |
| *Csf2ra* | 2.21E-07 | 1.386595 | 13.94559 | 0.000188 | 0.016439 |
| *Lrpap1* | 2.20E-07 | 1.386195 | 13.8968 | 0.000193 | 0.016575 |
| *Hint1* | 2.20E-07 | 1.386117 | 13.88731 | 0.000194 | 0.016575 |
| *Npc2* | 2.19E-07 | 1.384475 | 13.68888 | 0.000216 | 0.018217 |
| *Eef1g* | 2.16E-07 | 1.382032 | 13.39889 | 0.000252 | 0.021027 |
| *Atp5j* | 2.12E-07 | 1.377975 | 12.93016 | 0.000323 | 0.02631 |
| *Eif5a* | 2.12E-07 | 1.377958 | 12.9283 | 0.000324 | 0.02631 |
| *Gm2000* | 2.12E-07 | 1.377869 | 12.91813 | 0.000325 | 0.02631 |

| | | | | | |
|---|---|---|---|---|---|
| *Gabarap* | 2.12E-07 | 1.377256 | 12.84882 | 0.000338 | 0.027015 |
| *Dbi* | 2.11E-07 | 1.37636 | 12.74811 | 0.000356 | 0.028213 |
| *Gm8730* | 2.10E-07 | 1.374836 | 12.57848 | 0.00039 | 0.030573 |
| *Atp5h* | 2.09E-07 | 1.374561 | 12.54817 | 0.000397 | 0.030756 |
| *Ctsz* | 2.09E-07 | 1.374125 | 12.50022 | 0.000407 | 0.031237 |
| *Akr1a1* | 2.08E-07 | 1.373408 | 12.42164 | 0.000424 | 0.032253 |
| *Atp6v1f* | 2.07E-07 | 1.372371 | 12.30886 | 0.000451 | 0.033923 |
| *Uqcrb* | 2.07E-07 | 1.371708 | 12.23728 | 0.000468 | 0.034903 |
| *Ctsl* | 2.06E-07 | 1.37112 | 12.17421 | 0.000485 | 0.035753 |
| *Rap2b* | 2.06E-07 | 1.370421 | 12.09956 | 0.000504 | 0.036855 |
| *Syngr1* | 2.05E-07 | 1.369751 | 12.02838 | 0.000524 | 0.037925 |
| *Gpx4* | 2.05E-07 | 1.369448 | 11.99641 | 0.000533 | 0.038217 |
| *Gng5* | 2.04E-07 | 1.368978 | 11.94691 | 0.000547 | 0.038879 |
| *Gla* | 2.04E-07 | 1.368777 | 11.92576 | 0.000554 | 0.038885 |
| *Gm10076* | 2.04E-07 | 1.368647 | 11.91214 | 0.000558 | 0.038885 |
| *Ost4* | 2.03E-07 | 1.367772 | 11.82072 | 0.000586 | 0.04047 |
| *Ndufa13* | 2.03E-07 | 1.367411 | 11.7832 | 0.000598 | 0.040922 |
| *Cycs* | 2.02E-07 | 1.366411 | 11.67989 | 0.000632 | 0.042872 |
| *Cotl1* | 2.02E-07 | 1.366062 | 11.64399 | 0.000644 | 0.04332 |
| *Cox8a* | 2.01E-07 | 1.36587 | 11.62437 | 0.000651 | 0.043395 |
| *Selenow* | 2.01E-07 | 1.365009 | 11.5365 | 0.000682 | 0.044802 |
| *Romo1* | 2.01E-07 | 1.364972 | 11.5327 | 0.000684 | 0.044802 |
| *Pgam1* | 2.00E-07 | 1.36466 | 11.50109 | 0.000696 | 0.045181 |
| *Ms4a6c* | 2.00E-07 | 1.364437 | 11.47848 | 0.000704 | 0.045347 |
| *Anp32b* | 2.00E-07 | 1.364238 | 11.45836 | 0.000712 | 0.045455 |
| *Mmp12* | 1.99E-07 | 1.363302 | 11.36424 | 0.000749 | 0.047419 |
| *Ctsa* | 1.99E-07 | 1.362931 | 11.32709 | 0.000764 | 0.047645 |
| *Rbx1* | 1.99E-07 | 1.362768 | 11.31089 | 0.000771 | 0.047645 |
| *Uqcrh* | 1.99E-07 | 1.362755 | 11.30956 | 0.000771 | 0.047645 |
| *Atpif1* | 1.98E-07 | 1.36249 | 11.28312 | 0.000782 | 0.047938 |
| *Chchd10* | 1.98E-07 | 1.362115 | 11.24589 | 0.000798 | 0.04822 |
| *Tpi1* | 1.98E-07 | 1.362081 | 11.24255 | 0.000799 | 0.04822 |
| *Gm10053* | 1.97E-07 | 1.361375 | 11.17275 | 0.00083 | 0.049588 |
| *H2-D1* | 1.97E-07 | 1.36126 | 11.1614 | 0.000835 | 0.049588 |

#### Supplementary Table S4. 65 genes perturbed by the virtual-KO of *Hnf4a* and *Hnf4g* in intestinal cells.

The STRING interaction network of perturbed genes is available at the permalink: https://version-11-0b.string-db.org/cgi/network?networkId=bGGZHJwMOcny

| GENE | DISTANCE | Z | FC | P.VALUE | P.ADJ |
|---|---|---|---|---|---|
| *Hnf4g* | 3.23E-06 | 3.550427 | 126257.3 | 0 | 0 |
| *Hnf4a* | 1.16E-06 | 3.16971 | 16457.4 | 0 | 0 |
| *Ppia* | 1.21E-07 | 2.398759 | 176.2732 | 3.16E-40 | 2.63E-37 |
| *Eef1a1* | 9.65E-08 | 2.328612 | 113.0699 | 2.08E-26 | 1.30E-23 |
| *Tmsb10* | 9.39E-08 | 2.320066 | 107.074 | 4.29E-25 | 2.14E-22 |
| *Sepp1* | 8.91E-08 | 2.303584 | 96.36944 | 9.53E-23 | 3.97E-20 |
| *Hsp90ab1* | 8.84E-08 | 2.301246 | 94.93733 | 1.97E-22 | 7.01E-20 |
| *Fau* | 7.43E-08 | 2.246924 | 66.9273 | 2.82E-16 | 8.79E-14 |
| *Gpx1* | 7.02E-08 | 2.229493 | 59.7799 | 1.06E-14 | 2.94E-12 |
| *Apoa4* | 7.00E-08 | 2.228878 | 59.54163 | 1.20E-14 | 2.99E-12 |
| *Guca2b* | 6.76E-08 | 2.217887 | 55.438 | 9.65E-14 | 2.19E-11 |
| *Krt19* | 6.72E-08 | 2.215989 | 54.75783 | 1.36E-13 | 2.84E-11 |
| *Tomm7* | 6.59E-08 | 2.210323 | 52.77464 | 3.74E-13 | 7.18E-11 |
| *Apoc3* | 6.46E-08 | 2.203969 | 50.6341 | 1.11E-12 | 1.98E-10 |
| *Atp5j2* | 6.36E-08 | 2.19911 | 49.05397 | 2.49E-12 | 4.14E-10 |
| *Mdh1* | 6.34E-08 | 2.198147 | 48.74669 | 2.91E-12 | 4.54E-10 |
| *Ndufa7* | 6.24E-08 | 2.193365 | 47.24749 | 6.26E-12 | 9.19E-10 |
| *Dnase1* | 6.07E-08 | 2.184966 | 44.72241 | 2.27E-11 | 3.15E-09 |
| *Sectm1b* | 5.87E-08 | 2.174829 | 41.84896 | 9.86E-11 | 1.30E-08 |
| *H2-Q2* | 5.61E-08 | 2.160953 | 38.20458 | 6.37E-10 | 7.95E-08 |
| *Apoa1* | 5.56E-08 | 2.158112 | 37.49744 | 9.15E-10 | 1.09E-07 |
| *Tceb2* | 5.53E-08 | 2.156313 | 37.05613 | 1.15E-09 | 1.30E-07 |
| *Myl6* | 5.31E-08 | 2.144225 | 34.21961 | 4.92E-09 | 5.34E-07 |
| *Uqcr11* | 5.15E-08 | 2.134722 | 32.13864 | 1.44E-08 | 1.49E-06 |
| *Ace2* | 5.09E-08 | 2.131136 | 31.38587 | 2.12E-08 | 2.11E-06 |
| *Lct* | 4.96E-08 | 2.123741 | 29.88697 | 4.58E-08 | 4.40E-06 |
| *Smim24* | 4.78E-08 | 2.112376 | 27.71809 | 1.40E-07 | 1.30E-05 |
| *H2-Q1* | 4.74E-08 | 2.110163 | 27.3138 | 1.73E-07 | 1.54E-05 |
| *Cyp4v3* | 4.55E-08 | 2.097856 | 25.16849 | 5.25E-07 | 4.52E-05 |
| *H2-Ab1* | 4.53E-08 | 2.096252 | 24.90115 | 6.03E-07 | 5.02E-05 |
| *Atpif1* | 4.49E-08 | 2.093754 | 24.49042 | 7.47E-07 | 5.90E-05 |
| *Cd36* | 4.49E-08 | 2.093596 | 24.46459 | 7.57E-07 | 5.90E-05 |
| *Pls1* | 4.44E-08 | 2.090496 | 23.96452 | 9.81E-07 | 7.42E-05 |
| *Tpt1* | 4.36E-08 | 2.084588 | 23.03873 | 1.59E-06 | 0.000117 |
| *Slc5a1* | 4.18E-08 | 2.072152 | 21.20229 | 4.13E-06 | 0.000295 |
| *Oaz1* | 4.11E-08 | 2.067267 | 20.52043 | 5.90E-06 | 0.000409 |
| *Scp2* | 3.98E-08 | 2.057242 | 19.18713 | 1.19E-05 | 0.000799 |
| *Apob* | 3.93E-08 | 2.054132 | 18.79093 | 1.46E-05 | 0.000958 |
| *Atp5l* | 3.93E-08 | 2.053719 | 18.73888 | 1.50E-05 | 0.000959 |
| *Ggt1* | 3.87E-08 | 2.048843 | 18.13524 | 2.06E-05 | 0.001284 |
| *Eef2* | 3.79E-08 | 2.043319 | 17.47421 | 2.91E-05 | 0.001773 |
| *Gng5* | 3.76E-08 | 2.040387 | 17.13284 | 3.49E-05 | 0.002071 |
| *Cd74* | 3.62E-08 | 2.029321 | 15.90201 | 6.67E-05 | 0.003872 |
| *Atp5e* | 3.58E-08 | 2.026315 | 15.58283 | 7.90E-05 | 0.00448 |

| | | | | | |
|---|---|---|---|---|---|
| *Ubl5* | 3.48E-08 | 2.018047 | 14.73656 | 0.000124 | 0.006857 |
| *Dgat1* | 3.48E-08 | 2.017533 | 14.6854 | 0.000127 | 0.006893 |
| *Vil1* | 3.45E-08 | 2.014785 | 14.41509 | 0.000147 | 0.007787 |
| *Cox5b* | 3.41E-08 | 2.011781 | 14.12506 | 0.000171 | 0.008733 |
| *Atp5k* | 3.41E-08 | 2.011736 | 14.12076 | 0.000171 | 0.008733 |
| *Itm2b* | 3.33E-08 | 2.005014 | 13.49242 | 0.00024 | 0.011957 |
| *Uqcr10* | 3.33E-08 | 2.004208 | 13.4189 | 0.000249 | 0.012191 |
| *Uqcrq* | 3.31E-08 | 2.003185 | 13.3262 | 0.000262 | 0.012563 |
| *Anpep* | 3.28E-08 | 2.00034 | 13.07155 | 0.0003 | 0.01412 |
| *Muc13* | 3.25E-08 | 1.997174 | 12.79372 | 0.000348 | 0.016075 |
| *Ndufs6* | 3.24E-08 | 1.99656 | 12.74055 | 0.000358 | 0.016238 |
| *H2-Aa* | 3.23E-08 | 1.995571 | 12.65527 | 0.000375 | 0.016692 |
| *Neat1* | 3.21E-08 | 1.993702 | 12.49569 | 0.000408 | 0.017861 |
| *Atf3* | 3.18E-08 | 1.990885 | 12.25881 | 0.000463 | 0.019928 |
| *Ndufa12* | 3.16E-08 | 1.989503 | 12.14422 | 0.000492 | 0.020801 |
| *Ndufv3* | 3.16E-08 | 1.989156 | 12.11561 | 0.0005 | 0.020801 |
| *Cox17* | 3.12E-08 | 1.985689 | 11.83333 | 0.000582 | 0.023806 |
| *Mrpl12* | 3.09E-08 | 1.983051 | 11.62285 | 0.000651 | 0.026227 |
| *Myo15b* | 3.06E-08 | 1.979949 | 11.37997 | 0.000742 | 0.029413 |
| *Hspa8* | 3.04E-08 | 1.977454 | 11.18817 | 0.000823 | 0.032105 |
| *Uqcrb* | 2.94E-08 | 1.968472 | 10.52345 | 0.001179 | 0.045262 |

#### Supplementary Table S5. 17 genes perturbed by the virtual-KO of *Cftr* in alveolar type II cells.

The STRING interaction network of perturbed genes is available at the permalink: https://version-11-0b.string-db.org/cgi/network?networkId=bbh2WkHUgVGT

| GENE | DISTANCE | Z | FC | P.VALUE | P.ADJ |
|---|---|---|---|---|---|
| *Cftr* | 5.15E-06 | 4.919301912 | 52793.23173 | 0 | 0 |
| *Lamp3* | 1.00E-06 | 4.291962566 | 2004.709835 | 0 | 0 |
| *Sftpc* | 6.91E-07 | 4.156084723 | 951.5518206 | 6.10E-209 | 1.32E-205 |
| *Cxcl15* | 6.10E-07 | 4.110976075 | 740.7119933 | 4.20E-163 | 6.79E-160 |
| *Sftpa1* | 5.85E-07 | 4.096133987 | 681.8783452 | 2.61E-150 | 3.38E-147 |
| *Sftpb* | 5.72E-07 | 4.087956798 | 651.437397 | 1.09E-143 | 1.17E-140 |
| *Tspan1* | 3.82E-07 | 3.9446971 | 290.1749266 | 4.55E-65 | 4.21E-62 |
| *Birc5* | 3.08E-07 | 3.869868221 | 188.9493357 | 5.39E-43 | 4.36E-40 |
| *Pclaf* | 2.68E-07 | 3.821586141 | 142.908672 | 6.15E-33 | 4.43E-30 |
| *Dcxr* | 2.67E-07 | 3.820839046 | 142.2902594 | 8.40E-33 | 5.44E-30 |
| *Cldn10* | 2.63E-07 | 3.815853846 | 138.2300104 | 6.49E-32 | 3.82E-29 |
| *Smc2* | 2.61E-07 | 3.813323335 | 136.2124566 | 1.79E-31 | 9.67E-29 |
| *Hmgb2* | 2.42E-07 | 3.786508098 | 116.5215499 | 3.65E-27 | 1.82E-24 |
| *Pglyrp1* | 2.36E-07 | 3.778653331 | 111.2993896 | 5.09E-26 | 2.35E-23 |
| *Tubb5* | 2.13E-07 | 3.743372114 | 90.52418901 | 1.83E-21 | 7.89E-19 |
| *Mgst1* | 1.72E-07 | 3.670129577 | 58.7486313 | 1.79E-14 | 7.25E-12 |
| *Npc2* | 9.47E-08 | 3.472600178 | 17.87466051 | 2.36E-05 | 0.00898522 |

#### Supplementary Figures

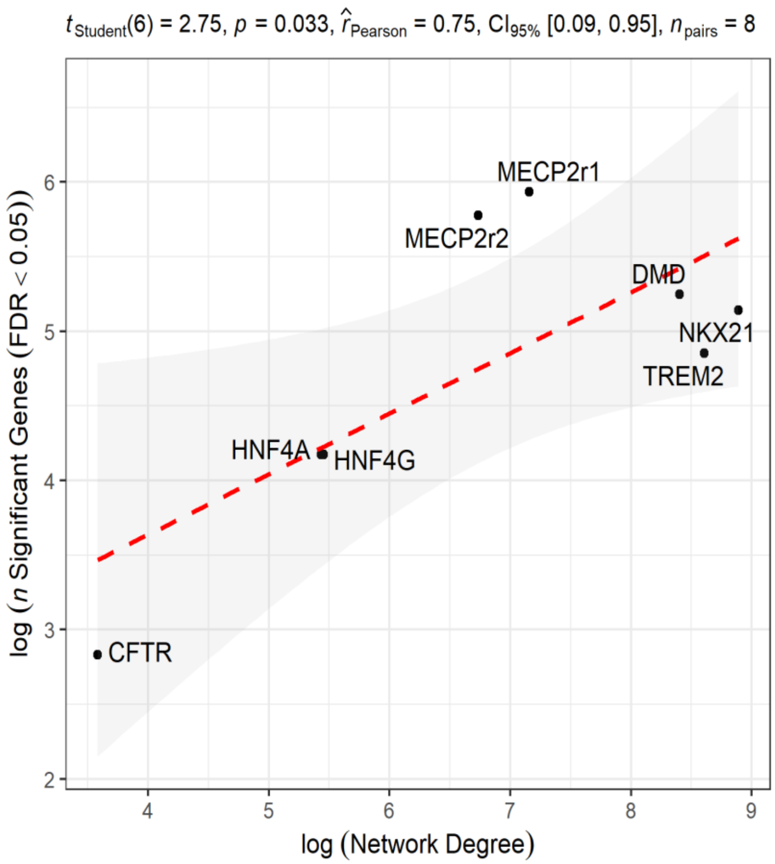

Supplementary Figure S1. Relationship between the number of significantly perturbed genes and the degree (i.e., the number of connections) of the KO gene in the network.

Note: x- and y-coordinates use a base of a natural logarithm scale on the x-axis and the y-axis

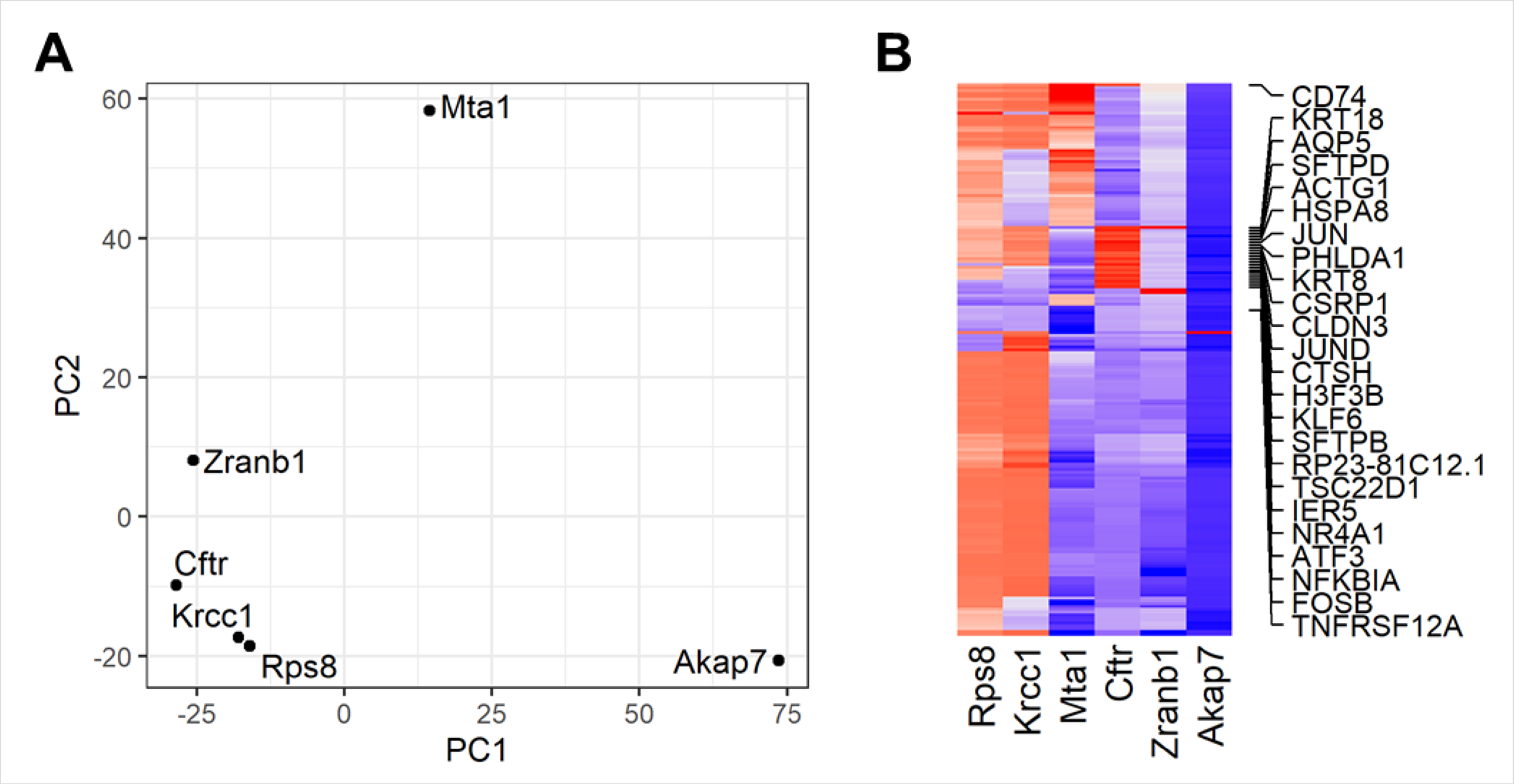

Supplementary Figure S2. Specificity analysis of the perturbation profiles provided by scTenifoldKnk in genes with similar expression patterns but different functionality.

**(A)** PCA of the perturbation profiles (ranked gene lists) generated by scTenifoldKnk for *Cftr* and 5 KO genes with similar expression profiles as *Cftr*. **(B)** Heatmap displaying the top 50 perturbed genes after the KO of genes.

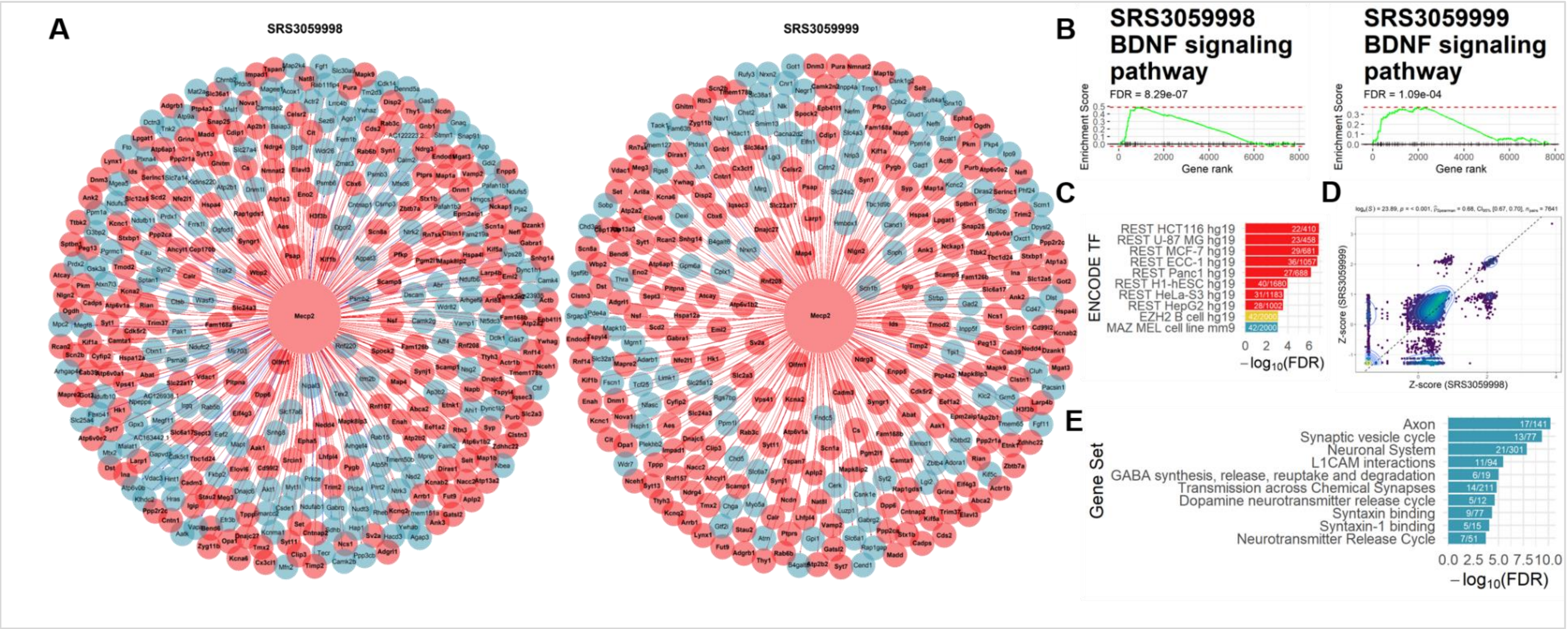

Supplementary Figure S3. scTenifoldKnk produces robust predictions using scRNAseq data from biological replicate samples. Identified predicted to be perturbed genes, transcription factors, and gene sets after Mecp2 gene knockout in mouse neurons. (**A**) Egocentric plots showing the identified predicted to be perturbed genes in two biological replicates. Genes are color-coded, in red if the gene is identified in both samples as significant (FDR < 0.05), and in blue, it was only identified as significant in one of the samples. (**B**) Gene set enrichment analysis showing association of the union of significant virtual-KO perturbed genes in the two replicates with the BDNF signaling pathway. (**C**) GSEA analysis shows the enrichment of the identified genes in both samples with the genes under the regulation of the REST transcription factor in 8 different cell lines. (**D**) Scatterplot showing the positive correlation between the Z-score transformed DR distances of genes, obtained using scTenifoldKnk Mecp2 KO analysis for two samples independently. (**E**) Gene set enrichment analysis showing the functional association of the predicted to be perturbed genes in both samples.

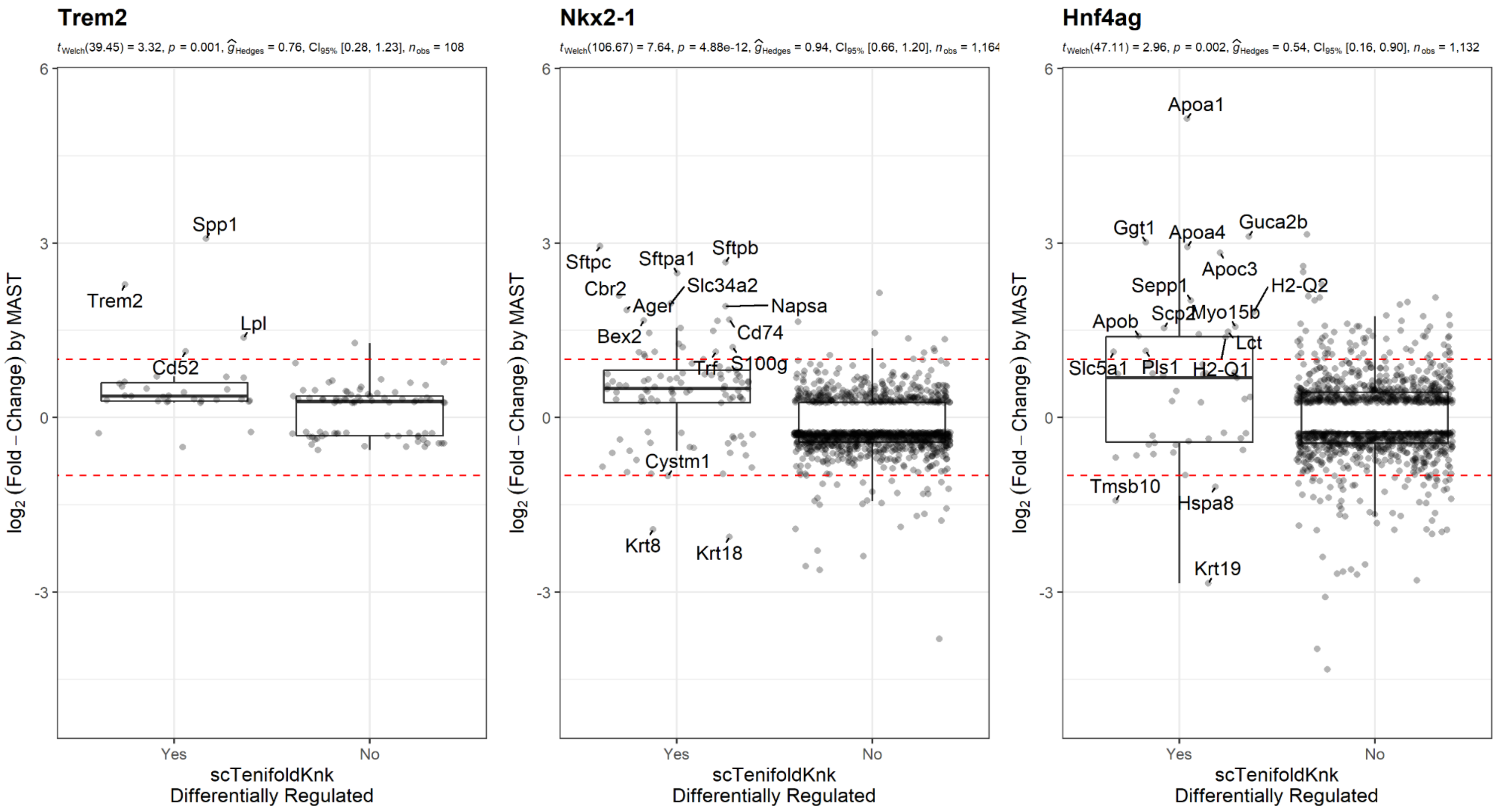

Supplementary Figure S4. Comparison of the values of fold-change (FC=WT/KO), as reported in differential expression analysis, between significantly perturbed genes and non-perturbed genes, as reported by scTenifoldKnk.

*Trem2* expression was detected in real-KO experiment and its expression was downregulated in KO samples (giving an increased FC value in the figure). Expression levels of *Nkx2-1* and Hnf4ag were too low in the KO samples to be included in DE analyses.

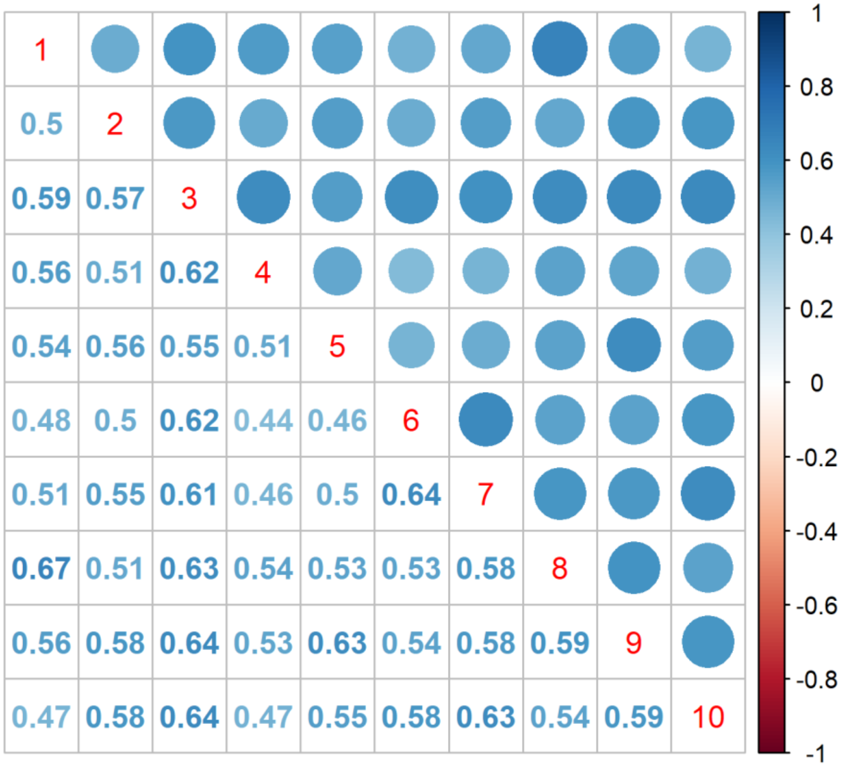

Supplementary Figure S5. Spearman correlation coefficients between perturbation profiles of 10 sets of subsampled Trem2 KO cells.

In the lower triangle, the values are reported numerically and in the upper triangle in a graphical manner. Dots are label proportional to the Spearman's correlation coefficients using the scale presented at the right of the matrix.

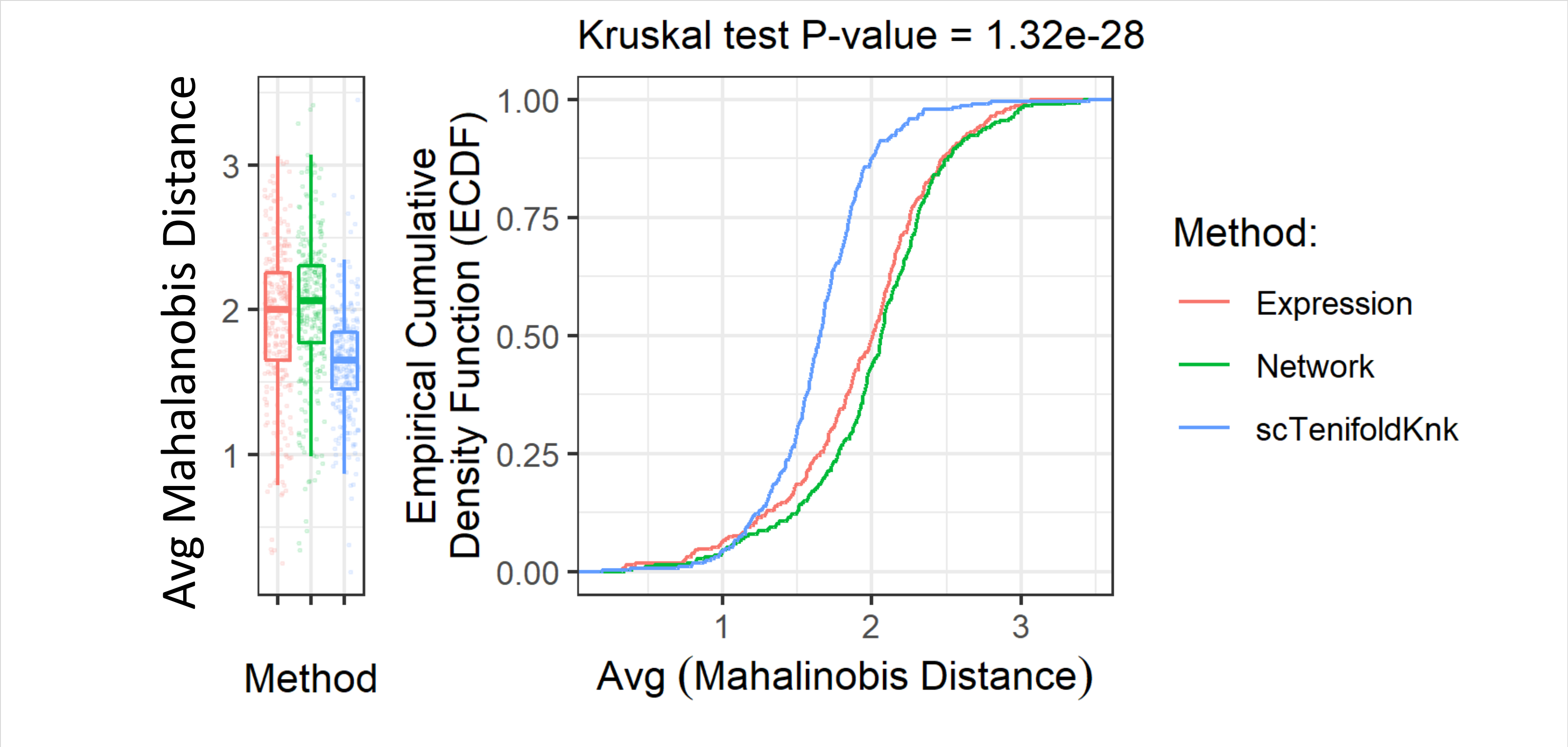

Supplementary Figure S6. Boxplot and CDF plot for the distribution of average Mahalanobis distance of genes to the centroid of given gene sets from the KEGG database.

The average distance from genes to their centroid in t-SNE embedding based on perturbation profile is significantly smaller than that based on expression profile and network edge weight profile. All 303 pathways included in the KEGG database (version 2019 available from Enrichr) were tested.

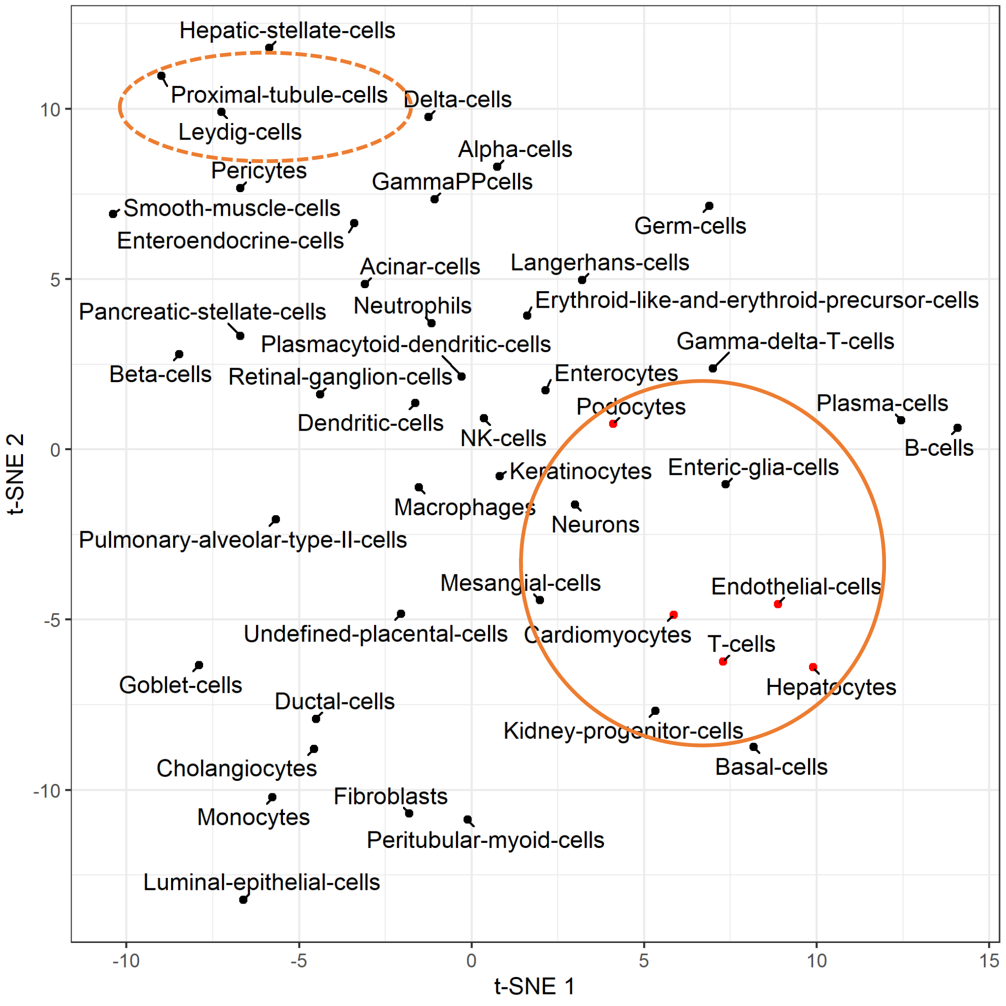

Supplementary Figure S7. tSNE representation of similarity and difference between cell types.

Virtual KO analysis was applied to all cell types presented here. The KO gene was *Mydgf*. The perturbation profiles of Mydgf obtained from different cell types were used as the input of tSNE. Cell types having similar profiles as endothelial cells are highlighted in red in the solid circle. Cell types mentioned in the main text, which are more different from endothelial cells, are shown in the dashed ellipse.
